# Systemic hypoxia inhibits T cell response by limiting mitobiogenesis via matrix substrate-level phosphorylation arrest

**DOI:** 10.1101/2020.03.19.999201

**Authors:** Amijai Saragovi, Ifat Abramovitch, Ibrahim Omar, Eliran Arbib, Ori Toker, Eyal Gottlieb, Michael Berger

## Abstract

Systemic oxygen restriction (SOR) is prevalent in numerous clinical conditions including chronic obstructive pulmonary disease (COPD). However, the influence of SOR on T cell protective immunity remains uncharacterized. Here we show the detrimental effect of hypoxia on mitochondrial biogenesis in activated CD8+ T cells. We find that low oxygen diminishes CD8+ T cell viral response *in vivo*. Using genetic and pharmacological models, we demonstrate that respiratory restriction inhibits ATP dependent matrix processes, all critical for mitochondrial biogenesis. The effect mediated by respiratory restriction could be rescued by TCA cycle re-stimulation, which led to increased mitochondrial matrix localized ATP via substrate-level phosphorylation. Finally, we demonstrate that short exposure to atmospheric oxygen pressure rescues the CD8+ viral response under systemic oxygen restriction *in vivo*. Our findings reveal the detrimental effect of hypoxia on mitochondrial biogenesis in activated CD8+ T cells and provide a new approach for reducing viral infections in COPD.

**Highlights:**

- Systemic chronic hypoxia compromises CD8+ T cell activation
- Shortly upon activation, T cells’ cytoplasmic activity becomes independent of mitochondrial ATP outflux
- Respiratory-blockade arrests mitochondrial remodeling due to energy depletion
- Uncoupler-based TCA stimulation rescues respiratory-restricted activated CD8+ T cells by stimulating matrix localized substrate-level phosphorylation
- CD8+ T cell arrest due to hypoxia *in vivo* can be rescued by short exposure to atmospheric oxygen pressure.

## INTRODUCTION

Our understanding of how systemic oxygen restriction (SOR) affects CD8+ T cell response remains incomplete. This is a difficult question to resolve as it requires the identification of specific metabolic effects within a dynamic system, that of activated cells in a process of rapid transformation and rewiring (MacIver et al., 2013). This is important as multiple respiratory and circulatory diseases including chronic obstructive pulmonary disease (COPD) and congenital heart disease are associated with hypoxemia, reduced blood oxygen saturation, and tissue hypoxia (Kent et al., 2011) (Kaskinen et al., 2016) (O’Brien and Smith, 1994). Notably, in numerous cases these patients were shown to have a higher prevalence of viral infections in comparison to healthy individuals (Chaw et al., 2019) (Kherad et al., 2010).

Previous studies have focused on the effect of hypoxia on cell fate determination in fully activated effector T cells (Doedens et al., 2013). They showed that hypoxic conditions contribute to the formation of long-lasting effector cells (Phan and Goldrath, 2015) (Phan et al., 2016). Other studies have revealed that respiratory restriction, mediated by inhibition of mitochondrial ATP synthase, arrests T cell activation (Chang et al., 2013). This is particularly interesting as activated T cells undergo a shift in cell metabolism early, in parallel to activation stimuli, switching to aerobic glycolysis to support their expansion and cytotoxic function (Gubser et al., 2013) (Van Der Windt et al., 2013). However, the mechanism leading to inhibition of T cell activation under hypoxic conditions remains uncharacterized.

In this study, we explore the effect of chronic systemic hypoxia on CD8+ T cell response. To assess the possible effect of systemic hypoxia *in vivo*, we first challenged mice with a lentivirus under conditions simulating COPD (Yu et al., 1999). We found that low oxygen availability diminished CD8+ T cell response to a lentivirus challenge. Similarly, hypoxic conditions applied *in vitro* led to complete arrest of CD8+ T cell response, but only marginally inhibited fully activated cells. To characterize the metabolic mechanism underlying T cell arrest, we chose to utilize the ATPsynthase inhibitor oligomycin, which allows distinguishing between indirect and direct effects of respiratory restriction. Incubation with oligomycin at different time points post stimuli revealed that following mitochondrial biogenesis, approximately twelve hours post stimuli, activated CD8+ T cells become independent of oxidative phosphorylation (OXPHOS). To understand why CD8+T cells are sensitive to respiratory restriction prior to mitochondrial biogenesis, we first examined cytoplasmic response to respiratory restriction via metabolic profiling and p-AMPK analysis. We found that respiratory restriction prior to mitochondrial biogenesis had a marginal effect on cytoplasmic function. In line with these results, inhibition of mitochondrial ATP transport to the cytoplasm by genetic alteration and pharmacological treatment had little effect on CD8+ T cell activation. In contrast, respiratory restriction prior to T cell mitochondrial biogenesis led to an energetic crisis within the mitochondrial matrix manifested by dysfunction in mitochondrial RNA processing and protein import. Application of a proton ionophore, FCCP, which uncouples the electron transport chain from ATPsynthase, rescued oligomycin-treated CD8+ T cells. Comparative metabolic profiling of activated T cells following uncoupler rescue, showed a significant increase in the generation of mitochondrial matrix-localized ATP via mitochondrial-localized substrate-level phosphorylation. Our findings establish that during early activation OXPHOS is required primarily to provide ATP for mitochondrial remodeling. Applying our insights to our *in vitro* model, we demonstrate that the detrimental effects of hypoxia may be alleviated by oxygen resuscitation.

## RESULTS

### Systemic chronic oxygen restriction inhibits CD8+ T cell activation and response

Clinical chronic hypoxia, prevalent in multiple respiratory and circulatory diseases, is associated with increased viral infections (Chaw et al., 2019) (Kherad et al., 2010). To test the effect of chronic hypoxia on CD8+ T cells’ response *in vivo*, we challenged mice with an OVA expressing lentivirus (L_OVA_) in a murine chronic hypoxia model. L_OVA_ challenge under low oxygen concentrations (8%) resulted in diminished CD8+ T cell response in respect to control, manifested by a substantially lower number of activated (CD62L negative), OVA associated (TCR Va2+), CD8+ T cells in the spleen and peripheral blood (Figures 1A-D). Our findings suggest that induction of systemic chronic hypoxia *in vivo* perpetuates the CD8+ T cell response.

**Figure 1:**
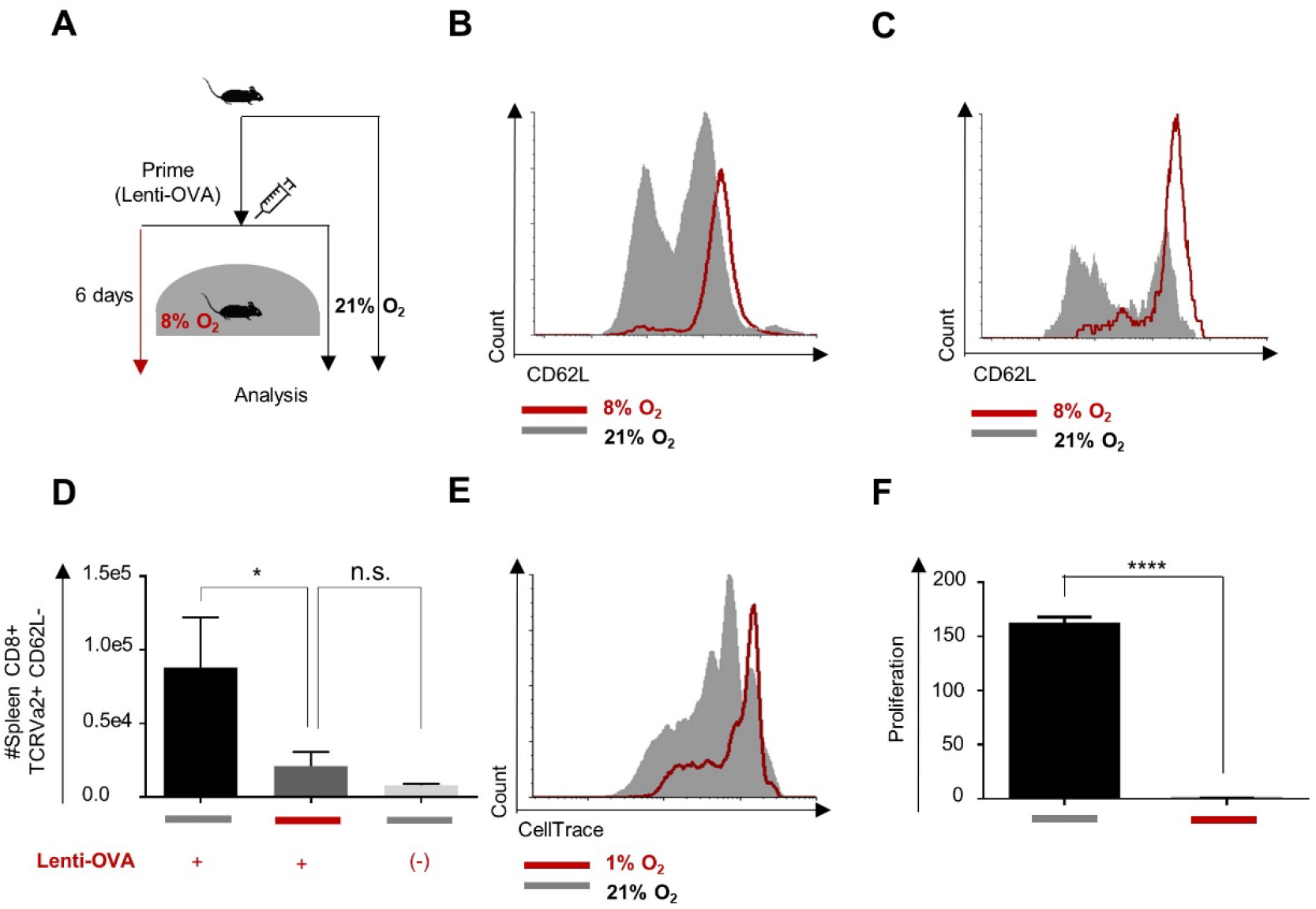
Systemic chronic oxygen restriction inhibits CD8+ T cell activation and response. **A** schematic of experiment presented in panel (B-D). OT1 splenocytes were adoptively transferred to WT mice, 2m splenocytes per mouse. Following 24 hours, mice were challenged with OVA expressing lentivirus or left untreated and transferred and then kept for 6 days under 8% or 21% oxygen pressure. **B** Representative histogram of CD62L expression in TCR Va2, CD8+ T cells extracted from spleens of mice challenged under 8% oxygen (red) or 21% oxygen (grey). **C** Representative histogram of CD62L expression in TCR Va2, CD8+ T cells extracted from peripheral blood of mice challenged under 8% oxygen (red) or 21% oxygen. **D** Summary of experiments results (n= 5 mice in each group). (E-F) Cell Trace-labeled splenocytes were activated with anti-CD3/28 for 72 hours under either 1% or 21% of oxygen. **E** Representative flow cytometry histogram plots of Cell Trace intensity gated on CD8 T cell population. **F** Bar graphs summarize flow cytometry analysis of proliferation of the CD8+ T cell population (n=5 biological replicates). (unpaired t-test, mean±s.e.m, ** p<0.01, *** p<0.001, **** p < 0.0001).

Effector T cells (T_eff_) evolved a specialized metabolic network to overcome oxygen deficiency at the site of inflammation (Doedens et al., 2013). To better understand the lack of T cell response observed, we probed the proliferative capacity of CD8+ T cells activated in an oxygen-deficient environment *in vitro*. In line with our findings and previous reports (Chang et al., 2013), naïve CD8+ T cells (T_n_) activated under hypoxic conditions demonstrated reduced proliferative capacity compared to control (Figures 1E-F). Thus, under systemic chronic respiratory restriction, CD8+ T cell activation is compromised.

### Following mitochondrial remodeling, activated CD8+ T cells become tolerant to inhibition of OXPHOS

To better understand the inhibitory effect mediated by respiratory restriction, we utilized the ATPsynthase specific inhibitor oligomycin, which partly mimics the effect induced by hypoxia (Chang et al., 2013) (Solaini et al., 2010) (Sgarbi et al., 2018). We chose to use oligomycin as it allows to distinguish between the indirect effects mediated by the inhibition of the electron transport chain and the TCA cycle (Martínez-Reyes et al., 2016) and the direct effect, as a result of reduced mitochondrial ATP (Lee and O’Brien, 2010). In a similar way to hypoxia, respiratory restriction mediated by an effective concentration (Figures S1A-C) of oligomycin led to inhibition of both mouse (Figure 2A) and human (Figures S1D-G) CD8+ T cell activation.

**Figure 2:**
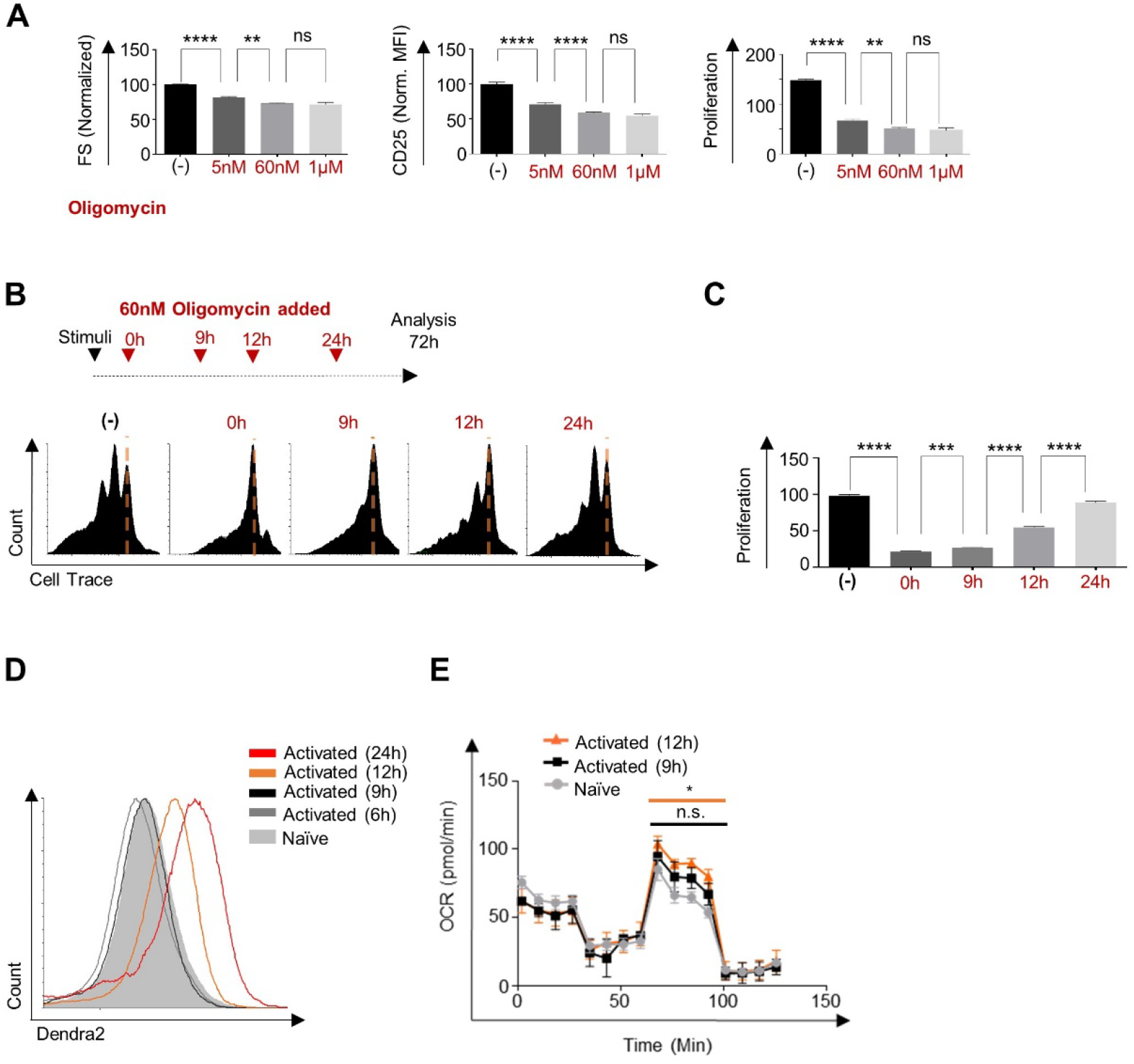
Following mitochondrial remodeling, activated CD8+ T cell become tolerant to inhibition of OXPHOS. **A** Cell Trace-labeled splenocytes were activated with anti-CD3/28 for 72 hours in the present or absent of indicated concentrations of or oligomycin. Bar graphs summarize flow cytometry analysis of cell size (left panels), CD25 expression levels (middle panels) and proliferation (right panels) of the CD8+ T cell population. n=5 biological replicates. **B** Cell Trace-labeled splenocytes were stimulated using anti-CD3/28 and treated with 60nM oligomycin at the indicated time points post activation. Seventy-two hours post activation, proliferation of CD8+ T cells were analyzed by flow cytometry. **C** Representative flow cytometry histogram plots of Cell Trace intensity gated on CD8 T cell population. n=5 biological replicates. **D** Flow cytometry histogram overlay plot of Dendra2 fluorescence intensity gated on the CD8+ T cell population from spleens of mito-Dendra2 of mito-Dendra2 mice that were stimulated using anti-CD3/CD28, for 6 (grey), 9 (black), 12 (orange), 24 (red) hours or left untreated (filled grey). n=6 biological replicates. **E** Mouse splenocytes were stimulated using anti-CD3/CD28 for 9 (black), 12 (orange) hours or left untreated (grey). CD8+ T cells were then isolated and assayed for OCR after consecutive injections of oligomycin, FCCP, and rotenone and antimycin (R+A) using seahorse XF24. n=5 biological replicates in each group. (unpaired t-test, mean ±s.e.m, * p<0.05, ** p<0.01, *** p<0.001, **** p < 0.0001).

To map the kinetics underlining the development of hypoxia tolerance, we examined CD8+ T cell response following oligomycin treatment at different time points post stimuli. Notably, when oligomycin was added at late activation, 12+ hours following stimuli (T_late_), we observed a substantial rescue in comparison to CD8+ T cells at early activation, up to 9 hours post stimuli (T_early_). T_late_ increased their cell size and demonstrated a marked increase in proliferation with respect to T_early_ following similar oligomycin treatment (Figures 2B-C, and S1H-K). Taken together, these observations suggest a gradual metabolic rewiring process, in which activated CD8+ T cells develop a metabolic bypass of respiratory restriction.

Glycolysis, the degradation of glucose to pyruvate/lactate, allow cells to generate ATP independent of oxygen concentrations (Lunt and Vander Heiden, 2011). Therefore, we tested whether the cellular capacity to perform glycolysis is correlated with the acquisition of tolerance to respiratory restriction during CD8+ T cell activation. To account for variances in glycolysis (Gubser et al., 2013) (Van Der Windt et al., 2013) extracellular acidification rate (ECAR) of T_n_, T_early_, and T_late_ were assessed using the seahorse methodology. Interestingly, both ECAR (oligomycin-untreated) and ECARmax (oligomycin-treated) were comparable in T_early_ and T_late_ and elevated in respect to T_n_ (Figure S2A-B, and S2C-E).

In line with the ECAR results, T_early_ demonstrated significant alterations in the concentration of key glycolysis-related metabolites with respect to T_n_ (Figure S2F). Specifically, T_early_ showed higher levels of intracellular glucose 6-phosphate (G6P), and dihydroxyacetone phosphate (DHAP) as well as a reduction in glyceraldehyde 3-phosphate (G3P) and glucose levels (Figure S2F). Similarly, secreted lactate and pyruvate concentrations were substantially elevated in T_early_ in comparison to T_n_ (Figure S2G). To test whether the rates of glucose uptake differ between early and late activation, we first incubated CD8+ T cells with the fluorescent glucose uptake probe, 2-deoxy-2-D-glucose (2-NBDG), at different time points following stimuli and analyzed their glucose uptake using flow cytometry. Supporting our previous findings, we detected no substantial difference in glucose uptake rate between T_n_, and CD8+ T cells activated for 6, 9 or 12 hours (Figure S2H). Noticeably, at a late stage of the activation, 24 hours following stimuli, we observed a considerable increase in the amount of glucose uptake. However, given the marked increase in cell volume, glucose concentrations are expected to remain the same (Figure S3 A-B). Collectively, these data demonstrate that stimulated T_n_ had a substantially elevated glycolytic rate at least 9 hours before they acquired tolerance to respiratory restriction. Thus, we find that the development of tolerance to respiratory restriction is not correlated with the increase in glycolytic activity.

In contrast to glycolysis, mitochondria were previously reported to undergo robust biogenesis and extensive metabolic rewiring ~12 hours post CD8+ activation (Rambold and Pearce, 2017) (Ron-Harel et al., 2016). To determine whether mitochondrial biogenesis correlates with the acquisition of tolerance to respiratory restriction, we measured the kinetics of mitochondrial biogenesis during CD8+ T cell activation in our model system. To this end, we first stimulated CD8+ T cells derived from mitochondria-labeled mtDendra2 mice (Pham et al., 2012). A substantial increase in mitochondrial mass was correlated with CD8+ T cell acquisition of tolerance to respiratory restriction (Figure 2D). Likewise, the maximum respiratory capacity significantly increased in late activation in respect to naive cells (Figure 2E). Thus, the development of tolerance to respiratory restriction during CD8+ T cell activation is correlated with gain of mitochondrial biomass linked to mitochondrial rewiring (Ron-Harel et al., 2016).

### Respiratory restriction has only a marginal effect on cytoplasmic function during early activation

CD8+ T cells can be activated following HIF1a stabilization (Doedens et al., 2013). In contrast, strong AMPK signaling may impair T cell activation (Araki et al., 2009) (Pearce et al., 2009). We therefore wanted to measure whether respiratory-restriction results in increased AMP-related signaling. To assess the effect of respiratory restriction on the level of different phospho-nucleotides, we first studied the metabolic profile of oligomycin-treated T_early_ and T_late_ with respect to untreated controls. As expected, oligomycin treatment led to a marked increase in mono/di-phospho-nucleotides at the expense of tri-phospho-nucleotides in T_early_ (Figure 3A). Hypoxia resilient T_late_ exposed to oligomycin demonstrated a similar effect (Figure 3B), suggesting that activated CD8+ T cells may function under increased AMP levels.

**Figure 3:**
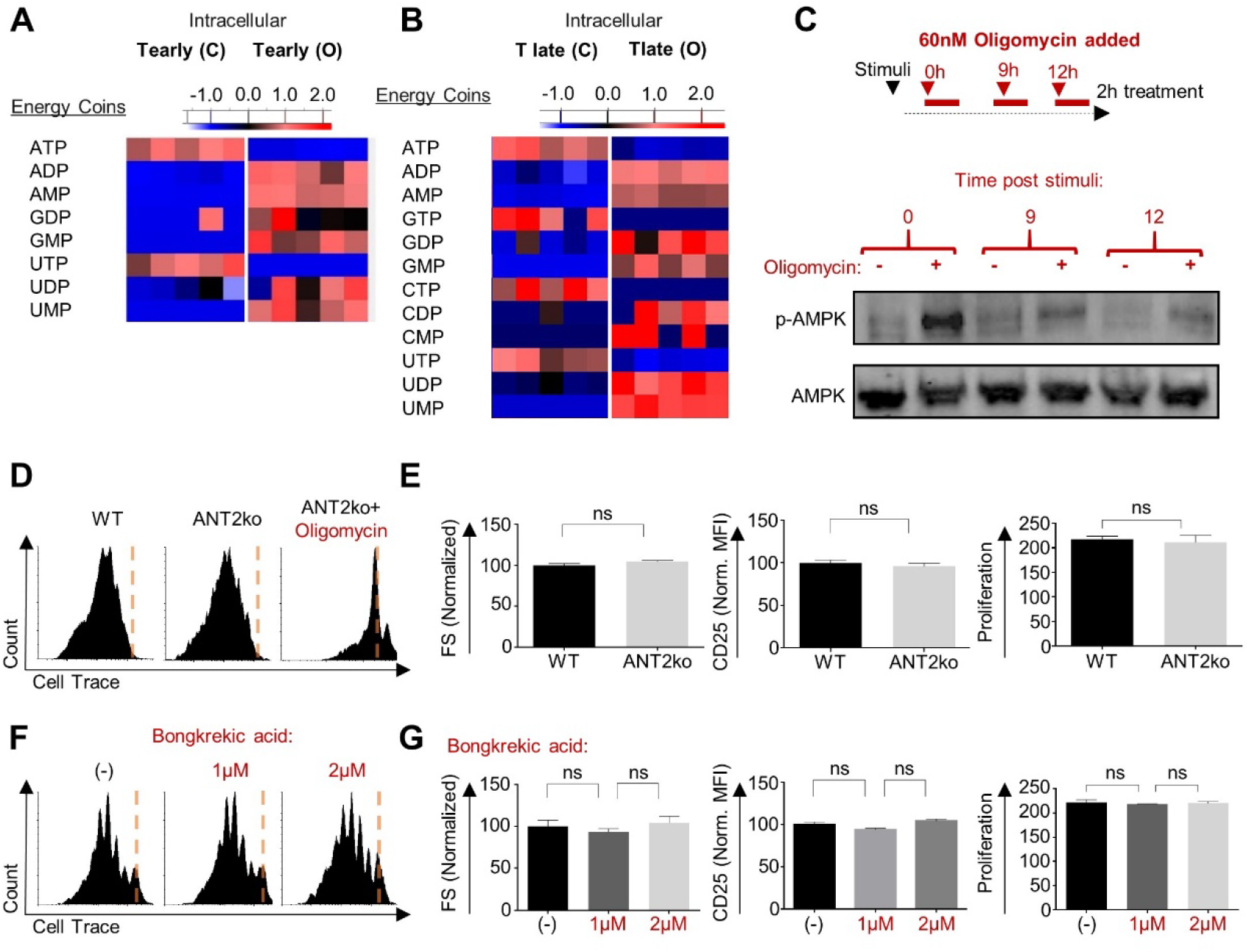
Respiratory restriction has only marginal effect on cytoplasmic function during early activation. **A-B** Heatmap showing relative amounts of key energy-related metabolites (as indicated in the figures) in oligomycin treated or untreated, CD8+ T cells activated for 5h (A) or for 24h (B). **C** Splenocytes were stimulated using anti-CD3/28 for 9 hours, 12 hours or left untreated (grey). Cells then were treated with 300nM oligomycin or left untreated for 1 hour. Protein extract from isolated CD8+ T cells from all samples were then subjected for immunoblot analysis using anti p-AMPK or anti AMPK. n=3 experiments. **D-E** Cell Trace-labeled splenocytes from wt or LCK-cre/ANT2floxp (ANT2ko) mice were stimulated using anti-CD3/CD28, with or without 60nM oligomycin. Seventy two hours post activation CD8+ T cells were analyzed for cell size, activation markers and proliferation by flow cytometry analysis. n=3 mice and 6 biological replicates. Representative flow cytometry histogram plots of Cell-Trace intensity gated on the CD8+ population (D). Bar graphs summarizing the experiment; cell size (left panel) CD25 expression (middle panel) and proliferation (right panel) of the CD8+ T cells population (E). **F-G** Cell Trace-labeled splenocytes from wt mice were stimulated using anti-CD3/CD28, in the presence of the indicated concentrations of the pan-ANT inhibitor, bongkrekic acid. Seventy two hours post activation, CD8+ T cells were analyzed for cell size, activation markers and proliferation by flow cytometry analysis. Representative flow cytometry histogram plots of Cell Trace intensity gated on CD8+ T cell population (F). Bar graphs summarize the cell size (left panel), CD25 expression (middle panel), and proliferation (right panel) of the CD8+ T cells in the experiment (G). n=6 biological replicates. (unpaired t-test, mean ±s.e.m).

Next, to evaluate the levels at which oligomycin-induced changes in phosphonucleotide profile affect cellular energy sensing, we examined the activation level of AMP-related signaling. To this end, we tested the phosphorylation level of AMP-activated protein kinase (p-AMPK, as a marker for AMPK activation), which is a cytoplasmic sensor for energy homeostasis, in CD8+ T cells activated for 9 and 12 hours. Strikingly, while oligomycin, the typical positive control for AMPK activation, markedly increased p-AMPK levels in treated naïve T cells, we observed only a marginal increase in p-AMPK following activation. Notably, the response to oligomycin was comparable in T_early_ and T_late_ (Figure 3C). Together, these results suggest that respiratory dependence during early T cell activation cannot be explained by alteration in the cellular response to reduced AMP levels in the cytoplasm.

Our findings so far imply that activated CD8+ T cells have a unique capacity to avoid p-AMPK signaling in the presence of elevated AMP levels. We therefore considered whether depleting mitochondrial ATP from CD8+ T cells’ cytoplasmic compartment will inhibit T_early_. For this purpose, we generated T cell-specific adenine nucleotide translocator 2 (ANT2, also called SLC25A5) knockout (ANT2ko) mice (Cho et al., 2017) (Cho et al., 2015). ANT2 is the dominant ADP/ATP translocator in murine CD8+ T cells, constituting approximately 90% of the total ANT protein (Figure S3A). Importantly, ANT2-ko CD8+ T cells displayed a substantial increase in mitochondrial membrane polarization (Figure S3B), which indicates a decrease in matrix ADP concentration.

To detect whether T cell-specific ANT2 deletion affects T cell activation, we examined the response to stimuli of ANT2-ko-derived CD8+ T cells. Surprisingly, ANT2-deficient T cells demonstrated normal activation-related phenotypes, including robust proliferative capacity (Figures 3D, 3E, and S3C). Noteworthy, ANT2-ko-derived CD8+ T cells were still sensitive to respiratory-restriction during early activation (Figures 3D, and S3C).

T cell-specific ANT2 deletion models chronic mitochondrial ATP restriction to the cytoplasm. To account for any compensatory effects that may have developed over time in these mice, and to observe the effect of acute mitochondrial ATP restriction to the cytoplasm, activated CD8+ T cells were treated with increasing doses of bongkrekic acid, a pan ANT inhibitor (Anwar et al., 2017). In line with our results, CD8+ T cells stimulated in the presence of bongkrekic acid at effective concentrations (Figure S3G), demonstrated normal activation and proliferation patterns (Figures 3F, 3G, and S3H). These key observations illustrate that ATP generated by mitochondrial respiration is not required for activated CD8+ T cell cytoplasmic function. Furthermore, these results point to an upstream respiratory-restriction-coupled effect as a limit underlying CD8+ T cells sensitivity to respiratory-restriction during the early stage of activation.

### Respiratory restriction leads to energetic crisis within the matrix compartment in early activated CD8+ T cells

Mitochondrial biogenesis and rewiring are critical checkpoints in T cell activation (Ron-Harel et al., 2016) (Rambold and Pearce, 2017). These cellular processes are dependent on the availability of matrix-bond ATP, which is generated by substrate-level phosphorylation, the metabolism of succinyl-CoA to succinate in the TCA cycle (Schwimmer et al., 2005) (Chinopoulos et al., 2010) depletes matrix-localized (Bochud-Allemann and Schneider, 2002). We therefore examined whether respiratory restriction ATP and affects mitochondrial biogenesis via an upstream effect. Oligomycin treatment at early CD8+ T cell activation abrogated the increase in mitochondrial mass observed in mtDendra2-derived CD8+ T cells (Figure 4A) suggesting that respiratory restriction disrupts mitochondrial biogenesis.

**Figure 4:**
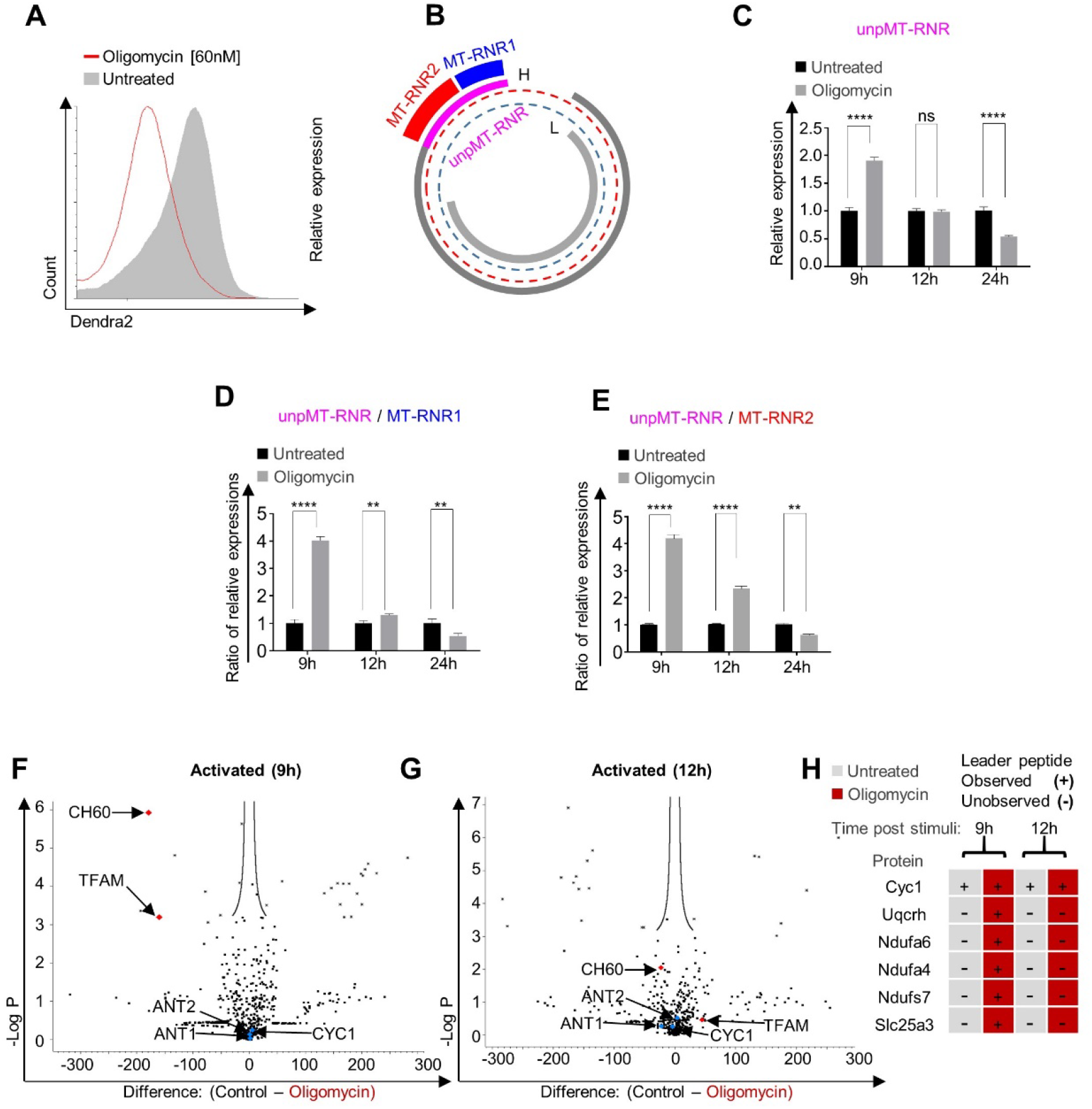
Respiratory restriction leads to energetic crisis within the matrix compartment in early activated CD8+ T cell. **A** Flow cytometry histogram overlay plot of Dendra2 fluorescence intensity gated on the CD8+ T cell population from spleens of mito-Dendra2 mice that were stimulated using anti-CD3/CD28 for 24 hours in the present or absent of 60nM oligomycin. n=3 independent experiments. **B** schematic of typical mitochondrial transcription, describing the transcription of polycistronic mtRNA (dark grey) from the mitochondrial DNA heavy strand (H, red dotted line) following by processing of the unprocessed MT-RNR transcript (unpMT-RNR, pink) to MT-RNR1 (bold blue line) and MT-RNR2 (bold red line). (C-E) Splenocytes were stimulated using anti CD3/CD28 for 9, 12 and 24 hours. One hour prior to CD8+ T cells isolation, cells were treated with oligomycin for one hour or left untreated. Total RNA then assayed using qRT-PCR for unprocessed MT-RNR transcript (unpMT-RNR) and the its two processed products, MT-RNR1 and MT-RNR2. **C** Relative expression of the unprocessed RNR. **D** Ratio between the relative expression of unpMT-RNR and MT-RNR1. **E** Ratio between the relative expression of unpMT-RNR and MT-RNR1 and 2, mitochondrial Encoded 12S and 16S rRNA respectively. n=6 biological replicates, 3 experiments (**F-H**) Splenocytes were stimulated using anti-CD3/CD28 for 9 or 12 hours. One hour prior to CD8+ T cells isolation, cells were treated with oligomycin. Protein extract from isolated CD8+ T cells were then subjected to immunoprecipitation using antiubiquitin antibody following by MS analysis focusing on mitochondrial proteins and mitochondrial leader peptides. n=3 biological replicates. Volcano plot of precipitated proteins detected by the MS analysis in oligomycin treated and untreated samples of 9 hours (**E**), or 12 hours (**F**). **G** comparison of leader peptides identified at 9 and 12 hours in respect to untreated samples. (unpaired t-test, mean ±s.e.m, *p<0.05 **p<0.01 *** p<0.001, **** p < 0.0001).

We next tested whether respiratory restriction will lead to an energetic crisis within the mitochondrial matrix. Taking a functional approach, we examined whether acute respiratory restriction disturbs two ATP-dependent processes within the matrix: protein import, in which mitochondrial-localized pre-proteins contain a leader sequence that is cleaved and removed upon entry to the matrix (Figure S4A) (Pfanner et al., 2019) (Wiedemann and Pfanner, 2017) (Chacinska et al., 2009); and processing of the polycistronic mitochondrial RNA coding for the 12S and 16S ribosomal RNAs (RNR1 and RNR2) (Wang et al., 2010) (Tu and Barrientos, 2015) (Buck et al., 2016). To examine whether respiratory restriction will lead to disturbance in matrix-localized RNA processing, we quantified both the relative expression levels of unprocessed RNR1-RNR2 polycistronic mitochondrial RNA (unpMT-RNR) and the ratio between unpMT-RNR and its processed products, RNR1 and RNR2 (Figure 4B) (Rackham et al., 2016). Oligomycin treatment led to a marked increase in the levels of unpMT-RNR in T_early_ but not in T_late_ (Figure 4C). In addition, and in line with these results, the ratios between unpMT-RNR and its cleaved products, RNR1 or RNR2, were found to be significantly further increased in oligomycin-treated T_early_ in comparison to T_late_ (Figures 4D-E). In a similar way to RNA processing, during matrix ATP deficiency, the import machinery is unable to pull nuclear-encoded matrix proteins. As a result of protein import disturbance, matrix and inner membrane proteins misfold in the cytoplasm leading to their degradation via the ubiquitin-proteasome pathway (Chacinska et al., 2009). We therefore assessed whether respiratory restriction will also lead to the accumulation of ubiquitinated mitochondrial matrix proteins and specifically ubiquitinated matrix pre-proteins. For this aim, T_late_ and T_early_ were treated with oligomycin for one hour or left untreated. Protein extracts from all samples were then subjected to immunoprecipitation using anti-ubiquitin antibody and assayed using mass spectrometry. As expected, acute oligomycin treatment significantly increased the amount of at least two central matrix proteins in T_early_ when compared to oligomycin-untreated controls. Specifically, T early had increased abundance of the mitochondrial transcription factor A, TFAM, and the CH60 chaperone, which plays a role in folding and assembly of newly imported proteins in the mitochondria. In line with the partial tolerance to oligomycin observed in late activation, we detected no substantial increase in matrix proteins in samples extracted from oligomycin-treated T_late_ in comparison to untreated controls (Figure 4F-G). Importantly, the amounts of mitochondrial proteins ANT2, ANT1, and CYC1, which do not require matrix ATP for mitochondrial localization, did not significantly differ in treated samples and controls in any of the groups. Furthermore, in samples from oligomycin-treated T_early_ we were able to detect leader peptides of several mitochondrial proteins in which mitochondrial import is dependent on matrix ATP (Figures 4H, S4B, and Supplementary Table 1). In contrast, in all untreated samples and late activated, oligomycin-treated samples, no relevant leader peptides were detected. Taken together, these results reveal a matrix-specific energetic crisis following respiratory restriction in early CD8+ T cell activation and point to TCA cycle inhibition and substrate-based-phosphorylation as a central inhibitory mechanism of respiratory restriction.

### FCCP treatment rescues respiratory-restricted CD8+ T cells by stimulating matrix-localized substrate level phosphorylation, elevates ATP and reduces AMP/GMP concentrations

TCA linked substrate-level phosphorylation is thought to fuel mitochondrial matrix activity, while ATP synthase derived ATP is exported to cytoplasm (Bochud-Allemann and Schneider, 2002) (Schwimmer et al., 2005). Oligomycin may indirectly lead to TCA cycle congestion, accumulation of intermediate metabolites and blockade of matrix-based substrate level phosphorylation. Thus, addition of uncouplers to respiratory ATP-deprived T_early_ cells may rescue the activation phenotype via TCA cycle stimulation (Figure S5 top). We therefore first attempted to rescue oligomycin treated CD8+ T cells by uncoupling the respiratory chain using an effective concentration of trifluoromethoxy carbonylcyanide phenylhydrazone (FCCP), a potent uncoupler of oxidative phosphorylation in mitochondria (Figures S5 bottom). T_early_ were treated with a fully effective oligomycin dose (Figures S1A-C), FCCP, both or left untreated. As expected, oligomycin treatment at early activation arrested CD8+ T cell proliferation. Notably, we observed that FCCP treatment alone slightly inhibited CD8+ T cell proliferation compared to control. Strikingly, the addition of FCCP to stimulated CD8+ T cells, which were treated with oligomycin, led to an almost complete rescue of proliferation compared to FCCP only treated cells (Figures 5A and 5B). These key observations demonstrate that uncoupling the respiratory chain from ATP synthase rescues respiratory restricted T_early_, suggesting that the maintenance of sufficient ATP concentrations in the matrix compartment is a key factor in the inhibitory mechanism that follows respiratory restriction.

**Figure 5:**
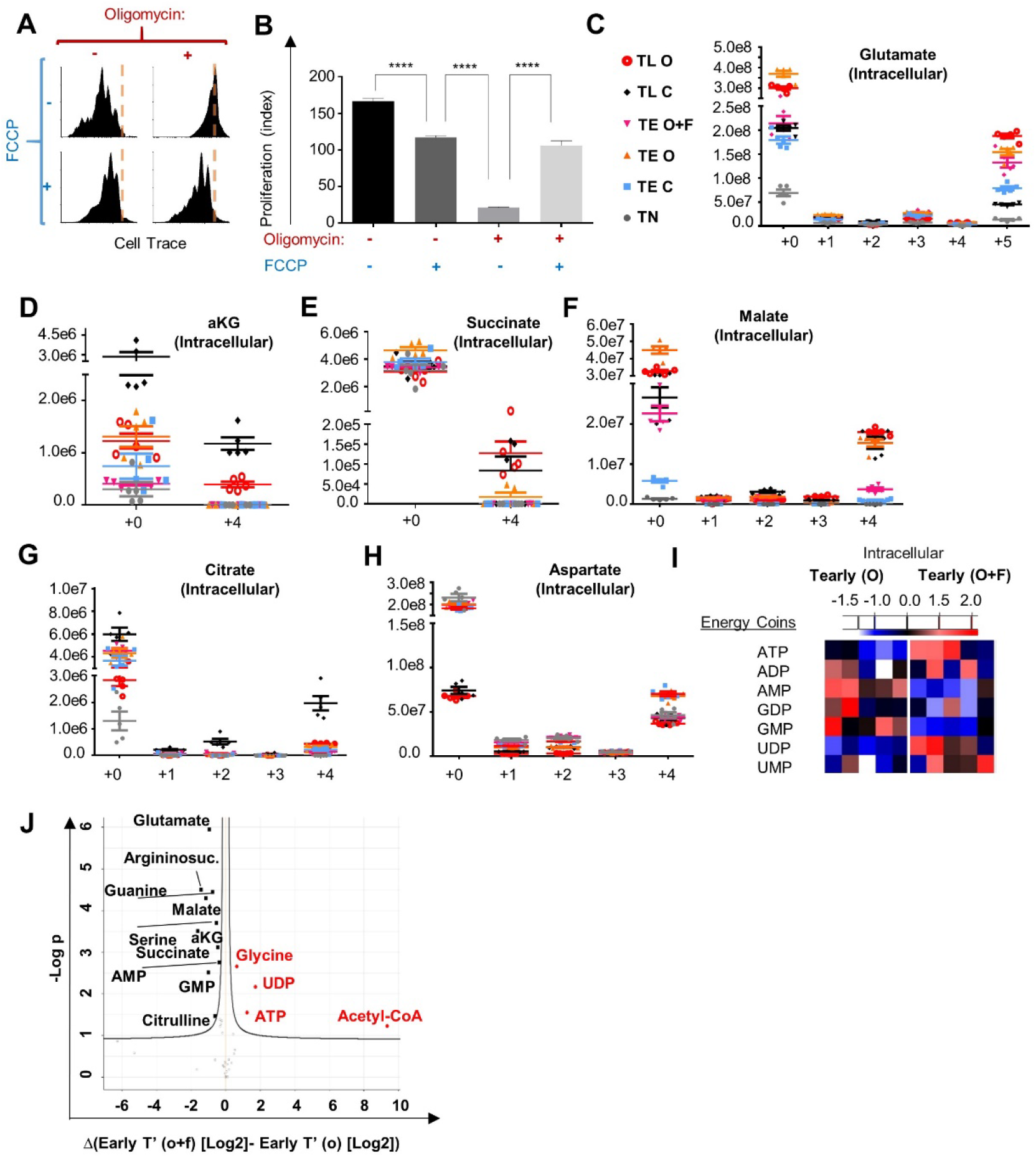
FCCP treatment rescues respiratory restricted CD8+ T cell by stimulating matrix localized substrate level phosphorylation, elevates ATP and reduces AMP/GMP concentrations. **A** Cell Trace-labeled splenocytes were stimulated using anti-CD3/28. Nine hours post activation cells were left untreated or treated with 1μM FCCP, 60nM oligomycin or combination of FCCP and oligomycin. Seventy-two hours post activation, proliferation of CD8+ T cells were analyzed by flow cytometry. Histograms show Cell Trace intensity of the CD8+ T cells population. n=5 biological replicates. **B** Bar graph summarizing the results presented in (A). (C-H) CD8+ T cells were activated with anti-CD3/CD28 for 5h, 9h or left unstimulated. Cells were then treated with media containing labeled glutamine with or without oligomycin, FCCP for 3 hours. Cells were then separately subjected for metabolomics analysis. T_late_ treated with oligomycin (TL O), T_late_ control (TL C), T_early_ treated with oligomycin + FCCP (TE O+F), T_early_ treated with oligomycin (TE O), T_early_ Control (TE C), Naïve (TN). Figures show **C** Bar graph summarizing intracellular isotopomers of glutamate, **D** aKG, **E** Succinate, **F** Malate, **G** Citrate, **H** Aspartate (n=7 biological replicates). **I** CD8+ T cells were activated with anti-CD3/CD28 for 5h. Cells then treated with oligomycin with or without FCCP for 3 hours. Cells were then separately subjected for metabolomics analysis. Figures show heat maps of relative amounts of energy related metabolites in the cells **J** Volcano plot of all the metabolites in T_early_ treated with oligomycin or oligomycin + FCCP.

To examine whether oligomycin treatment causes a surge in TCA cycle intermediates, we analyzed the metabolic profile of T_early_ incubated in media containing labeled glutamine following treatment with oligomycin or with both oligomycin and FCCP. As control we also examined a cohort of naïve CD8+ T cells, untreated T_early_, untreated T_late_, and oligomycin-treated T_late_. As can be seen in Figure 5C-G, respiratory restriction led to a marked increase in two key TCA cycle metabolites, glutamate and succinate, in both T_early_ and T_late_ compared to control. Interestingly, oligomycin treatment led to a marked increase in succinate (Figure 5E) and malate (Figure 5F) compared to control, but only at early activation. Notably in both T_early_, and T_late_, aspartate levels remain stable following oligomycin treatment (Figure 5H). Finally, we observed that the addition of FCCP to oligomycin treated T_early_ reduced the signal levels of all TCA-linked intermediates indicating stimulation of the TCA cycle (Figure 5C-G).

The release of the inhibition of the TCA cycle may allow respiratory-restricted cells to recover their matrix ATP via substrate-level phosphorylation, specifically via the conversion of succinyl-CoA to succinate. Accordingly, FCCP treatment is expected to allow respiratory restricted T_early_ to replenish their mitochondria with ATP, thus rescuing their matrix energy levels despite the inhibition of ATP synthase. To investigate this possibility, we looked at the levels of mono/di/tri-phosphonucleotides using metabolomics analysis. As expected, T_early_ treated with oligomycin and FCCP had significantly higher ATP and lower AMP/GMP concentrations compared to oligomycin only controls (Figures 5I-J). Given the ATP synthase blockade, the increase in cellular ATP and the reduction in both AMP and GMP may be attributed primarily to matrix-bound substrate-level phosphorylation.

### Short exposure to atmospheric oxygen pressure rescues CD8+ T cells’ response to lentiviral challenge under systemic hypoxia *in vivo*

Our findings demonstrate that during early activation, OXPHOS is required primarily to provide ATP for mitochondrial biogenesis. The buildup of additional mitochondrial biomass is thought to be a critical checkpoint in T cell activation (Ron-Harel et al., 2016) (Buck et al., 2016). Following the mitochondrial biogenesis checkpoint, fully activated CD8+ T cells show only marginal reduction in proliferation capacity under hypoxic conditions (Figure 6A) (Doedens et al., 2013). Thus, our results suggest that during systemic hypoxia, activated CD8+ T cells are arrested at the mitochondrial biogenesis checkpoint and may be rescued by short oxygen resuscitation. Building on these insights, we next attempted to rescue activated CD8+ T cell by re-exposing them to atmospheric oxygen following a period under hypoxic conditions (Figure 6B). Naive CD8+ T cells were activated under hypoxic or normal atmospheric conditions *in vitro*. After 24 hours, activated CD8+ T cells were re-exposed to normal atmospheric conditions or left under hypoxic conditions. As expected, activated CD8+ T cells that were left under hypoxic conditions for 72 hours were arrested and showed no elevation of activation markers (Figure 6C right panel) compared to control (Figure 6C left panel). In marked contrast, when CD8+ T cells activated under hypoxic conditions were re-exposed to atmospheric oxygen pressure, we observed a substantial increase in proliferative capacity (Figures 6C middle, 6D, and S6 A-B).

**Figure 6:**
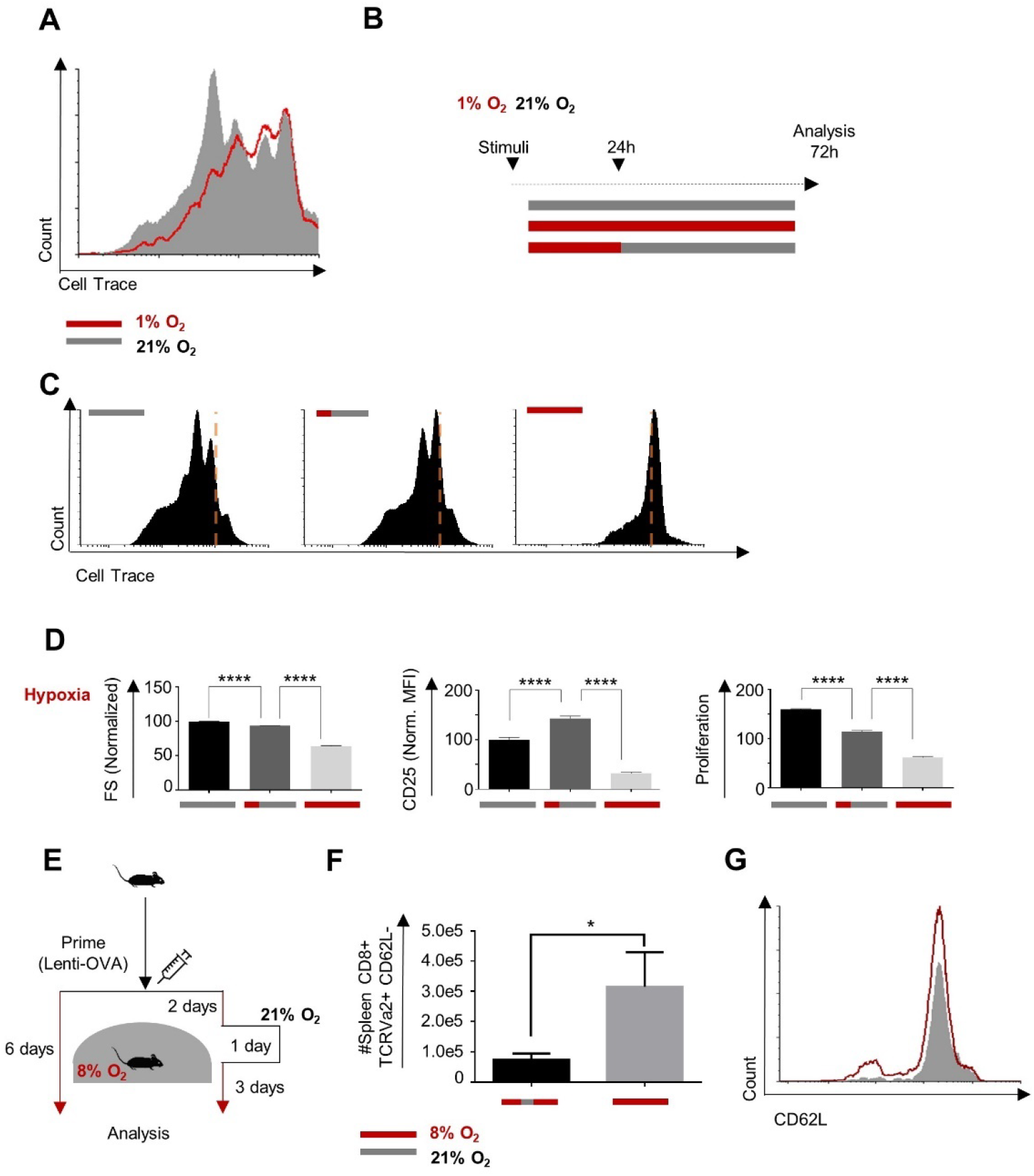
Short oxygen exposure rescues CD8+ T cells activated under hypoxia in vivo. **A** Cell Trace-labeled mouse splenocytes were stimulated using anti-CD3/28 for 24 hours. Cells were then transferred to hypoxia chamber containing 1% oxygen, or left untreated for additional 48 hours. Then, proliferation of the CD8+ T cells was analyzed using flow cytometry. **B** Schematic describing the experiment in (C) and (D). (C-D) Cell Trace-labeled splenocytes were stimulated with anti-CD3/28 for 72 hours, under normal conditions (top panels), hypoxic conditions (1% O2, 5% CO2, 94% N3) for the initial 24 hours and then in normoxic condition for 48 hours (middle panels), or under ongoing hypoxic conditions (bottom panels). CD8+ T cells were then analyzed by flow cytometry. **C** Representative histogram of Cell Trace staining. **D** Bar graphs summarizing the flow cytometry results presented in B-D. **F** Cell Trace-labeled splenocytes were activated with anti-CD3/28 for 72 hours in the present or absent of indicated concentrations of or oligomycin. Bar graphs summarize flow cytometry analysis of cell size (left panels), CD25 expression levels (middle panels) and proliferation (right panels) of the CD8+ T cell population. n=5 biological replicates. **E** Schematic of experiments presented in panel (F-G). OT1 splenocytes were adoptively transferred to WT mice, 2m splenocytes per mouse. Following 24 hours, mice were challenged with OVA expressing lentivirus and then kept for 6 days under 8% with or without a 24 hours resuscitation period in normal atmospheric conditions at day 3. **F** Summary of experiments results (n= 5 mice in each group). **G** Representative histogram of CD62L expression in TCR Va2, CD8+ T cells extracted from spleens of mice challenged under 8% oxygen with (red) or without (grey) 24 hours resuscitation period in normal atmospheric conditions at day 3. (unpaired t-test, mean ±s.e.m, * p<0.05).

Finally, to verify and extend our results, we attempted to rescue CD8+ response to lentivirus under systemic hypoxia by short exposure to atmospheric oxygen pressure *in vivo*. Mice were challenged with LOVA and then kept under 8% oxygen pressure for 6 days, with or without exposure for 24 hours to atmospheric oxygen pressure on day three (Figure 6E). In line with the *in vitro* results, short exposure to atmospheric oxygen pressure significantly improved the CD8+ T cell lentiviral response in mice kept under systemic hypoxia conditions (Figures 6F-G). Together, our results demonstrate that the detrimental effect caused by systemic hypoxia *in vivo* may be alleviated by short exposure to atmospheric oxygen pressure.

## DISCUSSION

In our study, we reveal that under systemic chronic hypoxia CD8+ T cell fail to activate and respond to a viral challenge due to matrix localized ATP deficiency which disturbed critical mitochondrial processes.

Several sets of evidence in our study indicate that mitochondrial respiratory based ATP is not required for T cell cytoplasmic function during activation. Using both genetic and acute models of cytoplasm-specific mitochondrial ATP restriction, we defined the demand for ATP in the mitochondrial matrix as a compartment distinct from the cytoplasm in CD8+ T cells. Furthermore, applying functional assays we were able to demonstrate a matrix-specific ATP crisis following acute respiratory-restriction. To confirm our results, we demonstrated that uncoupler-based restimulation of the TCA cycle can functionally rescue respiratory-restricted T_early_. In line with our findings, our metabolomic analysis revealed that oligomycin treatment at early activation led to an accumulation of TCA intermediates. Importantly we showed that addition of a proton uncoupler, FCCP, substantially reduced the accumulation of mono-phosphonucleotide intermediates and elevated ATP levels. Since oligomycin maintains ATPsynthase arrest even following the addition of FCCP (Lee and O’Brien, 2010), the increase in cellular ATP and the reduction in both AMP and GMP may be attributed primarily to matrix-bound substrate-level phosphorylation.

Our results suggest that during chronic hypoxia, activated CD8+ T cells are arrested at the mitochondrial biogenesis checkpoint and may be rescued by oxygen resuscitation. Building on our insights, we showed that *in vivo* hypoxia-arrested CD8+ T cells may be rescued by short exposure to atmospheric conditions.

Overall, our study revealed the detrimental effect of hypoxia on mitochondrial biogenesis in activated CD8+ T cells and provides a new approach to the reduction of viral infections in multiple hypoxia-associated diseases.

## EXPERIMENTAL PROCEDURES

### Mice

The C57BL/6J (wild-type), and Slc25a5tm1.1Nte/J *(ANT2^flox/lox^)* mice were from The Jackson Laboratory. B6.Cg-Tg(Lck-cre)1CwiN9 (Lck-Cre) were from Taconic. The T cell specific ANT2 knockout mice were generated by crossing mice containing a conditional floxed allele of ANT2 (Slc25a5tm1.1Nte/J *(ANT2flox/lox)* with transgenic mice expressing Cre under the control of the Lck gene promoter (Lck-Cre). Gt(ROSA)26Sortm1.1(CAG-Mito-Dendra2) Dcc (mito-Dendra2) mice were a kind gift from Dr. Tsvee Lapidot from the Weizmann Institute of Science. Mice were maintained and bred under specific pathogen free conditions in the Hebrew University animal facilities according to Institutional Animal Care and Use Committee regulations. All mice were maintained on the C57BL/6J background and used for experiments at 8–12 weeks of age.

### Quantitative Real-Time PCR and cDNA preparation

Total RNA from purified CD8+ T cells was extracted with Direct-zol RNA MiniPrep Plus (Zymo Research) following DNA removal step. cDNA was synthesized using ProtoScript First Strand cDNA Synthesis Kit (New England BioLabs, Inc – E6300L) with random primers for the MT-RNR transcripts, and oligo-dT primers for all other transcripts. Quantitative real-time PCR was then performed using Applied Biosystems (AB), Viia 7 Real Time PCR system with a Power SYBR green PCR master mix kit (Applied Biosystems).

Reaction was performed as follow:

1. 50°C 2 min, 1 cycle
2. 95°C 10 min, 1 cycle
3. 95 °C 15 s -> 60 °C 1 min, 40 cycles
4. 95 °C 15 s, 1 cycle
5. 60°C 1 min, 1 cycle
6. 95 °C 15 s, 1 cycle

Data was normalized to Mouse endogenous control (UBC and or RPL13) and analyzed using ΔΔCt model unless else is indicated.

Each experiment was performed in sixplicates and was repeated three times. Student’s t-test was used with 95% confidence interval.

### Primers used for quantitative Real-Time PCR

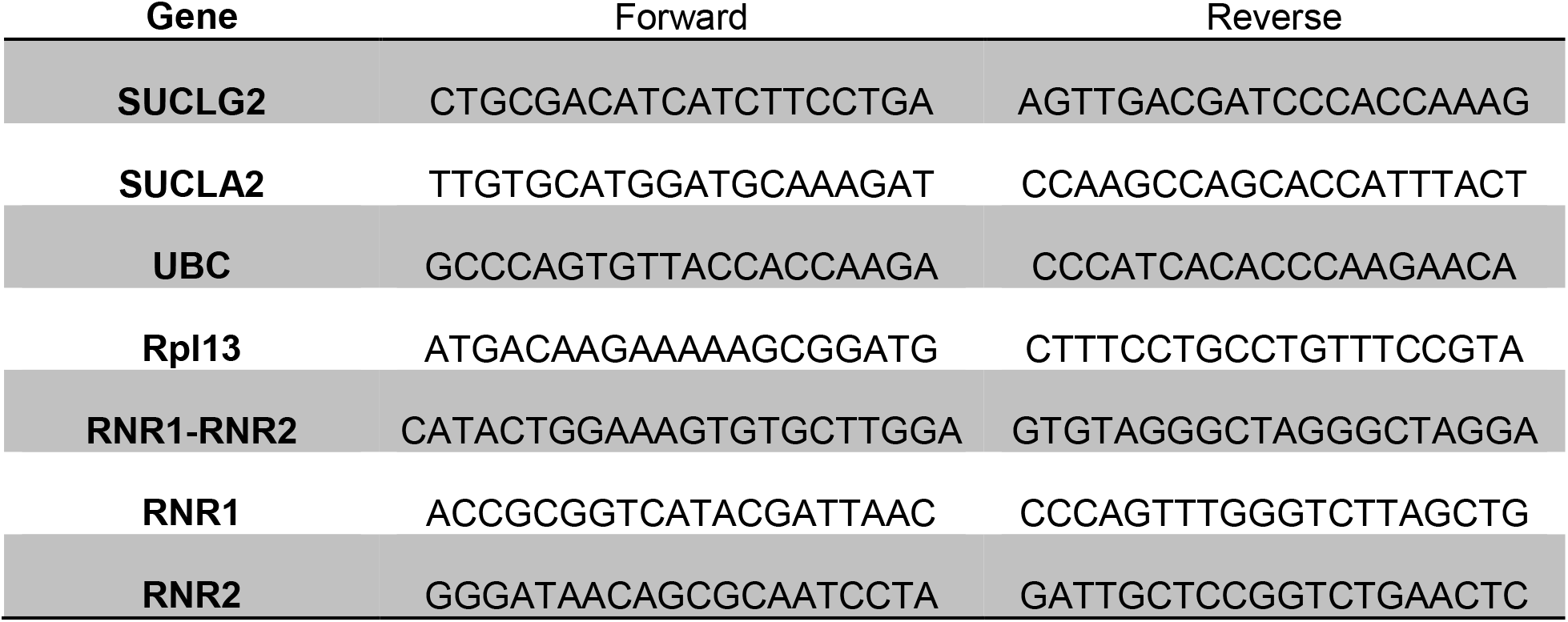

### Protein Mass Spectrometry

Sample preparation - Agarose beads containing immunoprecipitated samples, frozen at −20°C was subject to tryptic digestion, performed in the presence of 0.05% ProteaseMAX Surfactant (from Promega Corp., Madison, WI, USA). The peptides were then desalted on C18 Stage tips (Rappsilber et al., 2007). A total of 0.5 ug of peptides were injected into the mass spectrometer.

LC MS/MS analysis - MS analysis was performed using a Q Exactive Plus mass spectrometer (Thermo Fisher Scientific) coupled on-line to a nanoflow UHPLC instrument (Ultimate 3000 Dionex, Thermo Fisher Scientific). Eluted peptides were separated over a 60-min gradient run at a flow rate of 0.3 ul/min on a reverse phase 25-cm-long C18 column (75um ID, 2um, 100Å, Thermo PepMap®RSLC). The survey scans (380–2,000 m/z, target value 3E6 charges, maximum ion injection times 50 ms) were acquired and followed by higher energy collisional dissociation (HCD) based fragmentation (normalized collision energy 285). A resolution of 70,000 was used for survey scans and up to 15 dynamically chosen most abundant precursor ions were fragmented (isolation window 1.6 m/z). The MS/MS scans were acquired at a resolution of 17,500 (target value 5E4 charges, maximum ion injection times 57 ms). Dynamic exclusion was 60 sec.

MS data analysis - Mass spectra data were processed using the MaxQuant computational platform, version 1.5.3.12. Peak lists were searched against the Homo sapiens Uniprot FASTA sequence database containing a total of 26,199 reviewed entries or a custom FATSA file containing mouse mitochondrial leader peptides. The search included cysteine carbamidomethylation as a fixed modification and oxidation of methionine as variable modifications. Peptides with minimum of seven amino-acid length were considered and the required FDR was set to 1% at the peptide and protein level. Protein identification required at least 3 unique or razor peptides per protein group. The dependent-peptide and match-between-runs options were used.

### In vivo viral challenge under chronic hypoxia

OT1 splenocytes were adoptively transferred via intraperitoneal injection to WT mice (2 million splenocytes per mouse). Following 24 hours, mice were challenged with two intramuscular OVA expressing lentivirus injections (2.5* 10^6 TU suspended in 50ul per injection) or left untreated. Mice groups were then kept for 6 days under 8% or 21% oxygen pressure.

### *In vitro* T cell proliferation assay

Splenocytes or human PBMCs were stained with CellTrace (Molecular Probes, Eugene, OR) prior to activation for 30 min at 37°C. Cells were then activated in 24-flat-well plates (5 × 10^6^ cells per well) or 96-flat-well plates (1 × 10^6^ cells per well) with soluble anti-CD3ε (1μg/ml) and anti-CD28 (1μg/ml). For in-Seahorse activation of purified CD8+ T cells phorbol 12-myristate 13-acetate (PMA, 2nM) plus ionomycin (250ng/ml) were used.

### Antibodies

The following antibodies were used for flow cytometry: anti-CD8α (53-6.7), anti-CD4 (L3T4), anti-CD44 (IM7), CD122 (IL2RB) (5H4), CD127 (IL7R) (SB/199), anti-B220 (RA3-6B2), anti-CD69 (H1.2F3), anti-CD25 (3C7), anti-CD62L (MEL-14) and antihuman CD8 (HIT8a). All antibodies were from BioLegend.

Purified anti-CD3ε (145–2C11) and anti-CD28 (37.51; both from Biolegend) were used at the appropriate concentration for mouse T cell activation. Purified anti-CD3ε (OKT3) and anti-CD28 (CD28.2; both from Biolegend) were used at the appropriate concentration for human T cell activation.

Antibody to AMPKa phosphorylated on Thr172 and anti-AMPKa both from Cell Signaling Technology were used for immunoblot analysis.

Antibody to Ubiquitinated proteins (FK2) from Merck-Millipore was used for the immunoprecipitation assay.

### Flow Cytometry

Cells were stained with various conjugated mAbs against cell-surface markers in FACS buffer (PBS containing 1% FBS) for 30 min at 4°C. For mitochondrial membrane potential staining, cells were labeled with TMRM 50 nM (Molecular Probes, Eugene, OR) in FACS buffer without EDTA for 30 min at 30°C. Stained cells were analysed by Gallios flow cytometer with Kaluza software (Beckman Coulter, Brea, CA) and analysed by FACS Express 6 (De Novo Software).

### Metabolism Assays

OCR and ECAR were measured using a 24-well XF extracellular flux analyzer (EFA) (Seahorse Bioscience). Purified naive or activated CD8+ T cells (1 × 10^6^ cells per well) were seeded in Seahorse XF24 designated plates using Cell-Tak (Corning) adherent and assayed according to manufacturer instructions. In-Seahorse stimuli was performed using PMA (2nM) and Ionomycin (250ng/ml).

### Western Blot and Immunoprecipitation

Purified naive or activated CD8+ T cells were lysed in radioimmunoprecipitation assay (RIPA) buffer; 10 μg protein from each sample was separated by SDS–PAGE, and immunoblotted with with anti AMPK*α* antibody or p-AMPKa (Thr172) (Cell Signalling, Danvers, MA; 2532) followed by peroxidase donkey anti-rabbit IgG (Jackson Laboratory; 711-005-152).

For immunoprecipitation extracts from purified activated CD8+ T cells were prepared in extraction buffer (50 mM Tris-HCI, pH 8.0, 5 mm EDTA, 150 mM NaCl and 0.5% NP-40, supplemented with Protease Inhibitor Cocktail, Sigma-Aldrich, Israel). Protein extracts were then precleared with protein G beads (EZview™ Red Protein G Affinity Gel, Sigma-Aldrich, Israel), following incubation for 30 min at 4°C. Protein G beads were pelleted out, and the supernatant was taken for immunoprecipitation with 2 μg of anti-ubiquitin antibody (FK2, Merck-Millipore) for 12 hours at 4°C. Immune complexes were pelleted with protein G beads as before, and the pellets were washed three times in buffer B (5% sucrose, 50 mM Tris-HCI pH 7.4, 500 mM NaCI, 5 mM EDTA and 0.5% NP-40), followed by three washes with buffer C (50 mM Tris-HCI pH 7.4, 150 mM NaCl and 5 mM EDTA). The precipitated proteins were then subjected to MS analysis.

### Targeted metabolic analysis

CD8+ T cells were cultured in either anti-CD3/CD28 coated or uncoated 96 well plate (1 million cells /well), suspended in RPMI supplemented with 10% dialyzed Fetal Bovine Serum and 100μM Alanine with or without labeled glutamine. Following 5 or 24 hours activated cells were treated with 500nM Oligomycin, Oligomycin and 1μM FCCP or left untreated for 2 hours. Naïve, and activated cells were then extracted for metabolomics LC-MS analysis.

Medium extracts: Twenty microliters of culture medium was added to 980 μl of a cold extraction solution (−20°C) composed of methanol, acetonitrile, and water (5:3:2). Cell extracts: Cells were rapidly washed 3 times with ice-cold PBS, after which intracellular metabolites were extracted with 100μl of ice-cold extraction solution for 5 min at 4°C. Medium and cell extracts were centrifuged (10 min at 16,000g) to remove insoluble material, and the supernatant was collected for LC-MS analysis. Metabolomics data was normalized to protein concentrations using a modified Lowry protein assay.

LC-MS metabolomics analysis was performed as described previously (MacKay et al., 2015). Briefly, Thermo Ultimate 3000 high-performance liquid chromatography (HPLC) system coupled to Q-Exactive Orbitrap Mass Spectrometer (Thermo Fisher Scientific) was used with a resolution of 35,000 at 200 mass/charge ratio (m/z), electrospray ionization, and polarity switching mode to enable both positive and negative ions across a mass range of 67 to 1000 m/z. HPLC setup consisted ZIC-pHILIC column (SeQuant; 150 mm x 2.1 mm, 5 μm; Merck), with a ZIC-pHILIC guard column (SeQuant; 20 mm x 2.1 mm). 5 μl of Biological extracts were injected and the compounds were separated with mobile phase gradient of 15 min, starting at 20% aqueous (20 mM ammonium carbonate adjusted to pH.2 with 0.1% of 25% ammonium hydroxide) and 80% organic (acetonitrile) and terminated with 20% acetonitrile. Flow rate and column temperature were maintained at 0.2 ml/min and 45°C, respectively, for a total run time of 27 min. All metabolites were detected using mass accuracy below 5 ppm. Thermo Xcalibur was used for data acquisition. TraceFinder 4.1 was used for analysis. Peak areas of metabolites were determined by using the exact mass of the singly charged ions. The retention time of metabolites was predetermined on the pHILIC column by analyzing an in-house mass spectrometry metabolite library that was built by running commercially available standards.

### Statistical analysis

The statistical significance of differences was determined by the two-tailed Student’s *t*-test. Biological replicates refer to independent experimental replicates sourced from different mice/human donors. Technical replicates refer to independent experimental replicates from the same biological source. Differences with a *P* value of less than 0.05 were considered statistically significant. Graph prism and Perseus programs were use. MS data was normalized by ranking, when applicable, non-values were plugged with replicates mean to prevent zeros bias.

## ACKNOWLEDGMENTS

The authors thank Dr. Ofer Mandelboim (The Hebrew University) and Dr. David Chan (Caltech) for critical reviews, and Dr. Tsvee Lapidot (Weizmann Institute) for mito-Dendra2 mice.

This work was supported by grants from the ISRAEL SCIENCE FOUNDATION grant No. 1596/17, and in part by the German Israeli Foundation for Scientific Research and Development number I-1474-414.13/2018

## AUTHOR CONTRIBUTIONS

A.S. designed, performed research, analyzed data and wrote the manuscript; M.B., designed research, analyzed data and wrote the manuscript; I.O., E.A. and I.A. performed research; E.G. designed research and analyzed data; O.T. collected data

## SUPPLEMENTARY FIGURES

**Figure S1:**
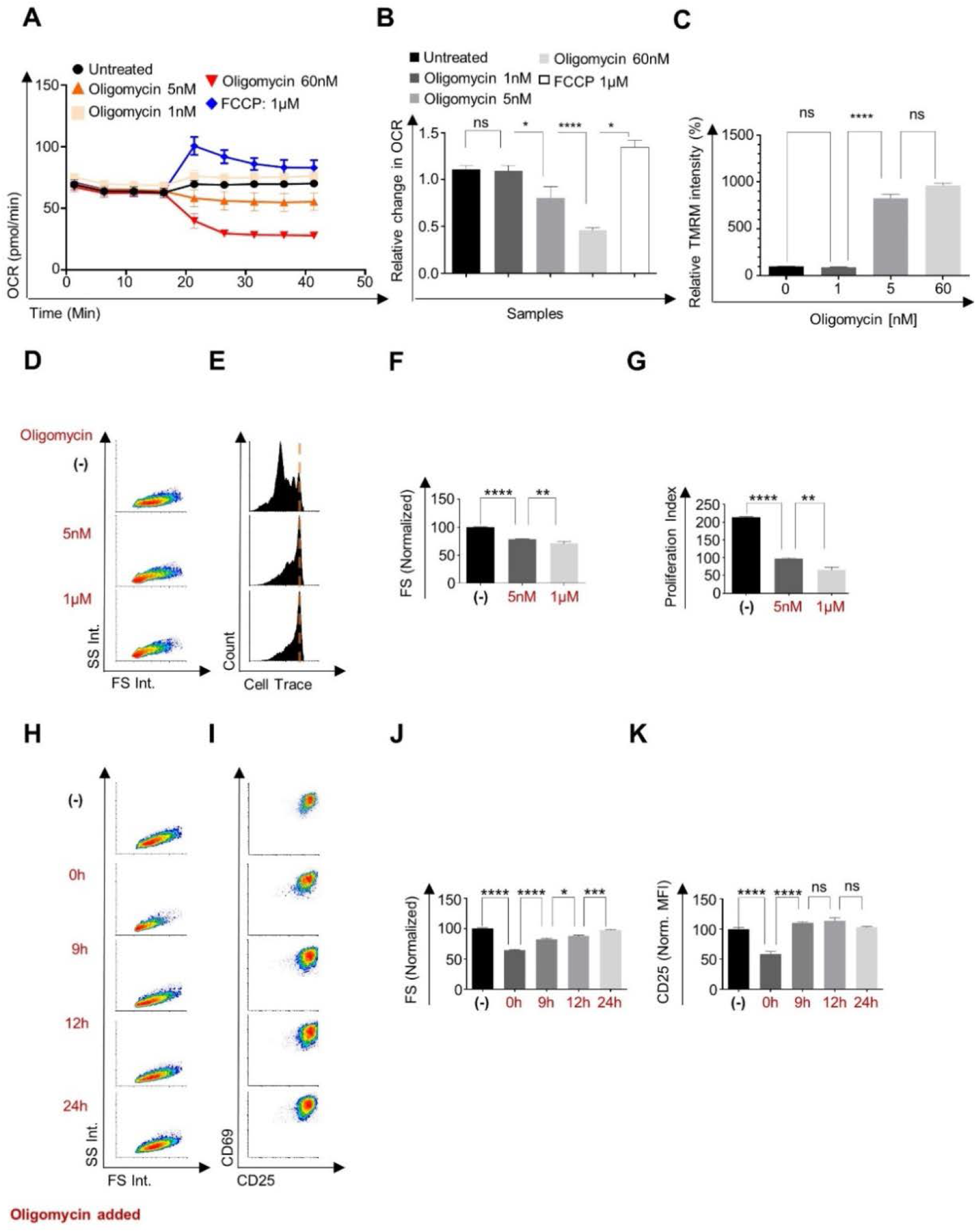
**A** OCR of isolated naive CD8+ T cells treated with the indicated oligomycin or FCCP concentrations. n=5 biological replicates. **B** Bar graph summarizes the relative change in OCR following oligomycin/FCCP treatment of the results shown in (A). C Bar graph summarizing the flow cytometry analysis of TMRM staining of naive CD8+ T cells treated with the indicated oligomycin concentrations. n=6 biological replicates. (D-E) Cell Trace-labeled human peripheral blood cells were stimulated with anti-CD3/28 for 72 hours in the present or absent of indicated concentrations of oligomycin. CD8+ T cells were then analyzed by flow cytometry for **D, F** size and granularity, **E, G** and proliferation. n=5 biological replicates. (**H-K**) Cell Trace-labeled mouse splenocytes were stimulated using anti-CD3/CD28 and treated with 60nM oligomycin at the indicated time points post activation. Seventy-two hours post activation, CD8+ T cells were analyzed by flow cytometry. n=5 biological replicates. **H** Representative flow cytometry plots of size and granularity, and **I** CD25 and CD69 expression levels gated on CD8+ T cell population. **J** Bar graphs summarize flow cytometry analysis of cell size (H), and CD25 expression levels (I). (unpaired t-test, mean±s.e.m, *p<0.05 **p<0.01 *** p<0.001, **** p < 0.0001).

**Figure S2:**
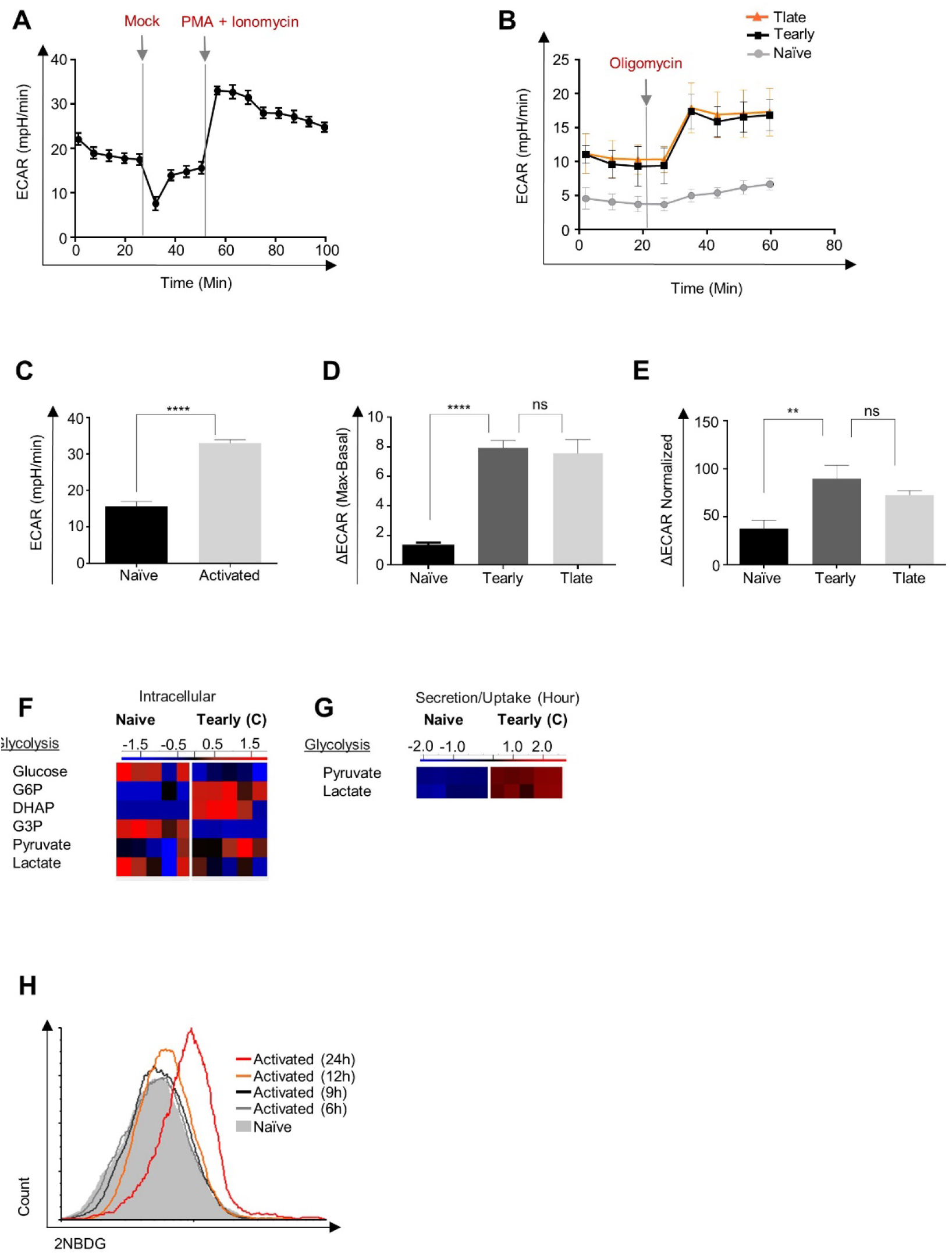
**A** Extracellular acidification rate (ECAR) measured by seahorse XF24 of isolated CD8+ T cells that were stimulated in-seahorse using PMA + Ionomycin. n=7 biological replicates. **B** Splenocytes were stimulated using anti-CD3/CD28 for 12 hours (orange), 9 hours (black) or left untreated (grey). CD8+ T cells were then isolated and assayed to quantify both basal and oligomycin induced maximum ECAR using seahorse XF24. n=5 biological replicates. **C** Bar graph summarizing (A) the extracellular acidification rate (ECAR) measured by seahorse XF24 of isolated CD8+ T cells that were stimulated in-seahorse using PMA + Ionomycin. **D** Bar graph presenting the delta between maximal ECAR (following oligomycin treatment) and basal ECAR of naive CD8+ T cells and activated CD8+ T cells for 9 or 12 hours. **E** Bar graph presenting normalized delta ECAR; (delta ECAR/Basal ECAR) x 100, of naive CD8+ T cells and activated CD8+ T cells for 9 or 12 hours. **F** Heatmap showing relative amounts of key glycolysis-related metabolites in naïve and activated for 5h CD8+ T cells. The strength of the color refers to how strongly upregulated (red) or downregulated (blue) the various metabolites are. **G** Heatmap showing relative amounts of pyruvate and lactate in the medium from naïve or activated for 5h (early activated CD8+ T cells. **H** Histogram overlay of 2NBDG fluorescence intensity gated on CD8+ T cells stimulated with anti CD3/CD28 for 6 (grey), 9 (black), 12 (orange), 24 (red) hours or unstimulated (filled-grey) that were incubated with 2NBDG for 30 min before flow cytometry analysis.

**Figure S3:**
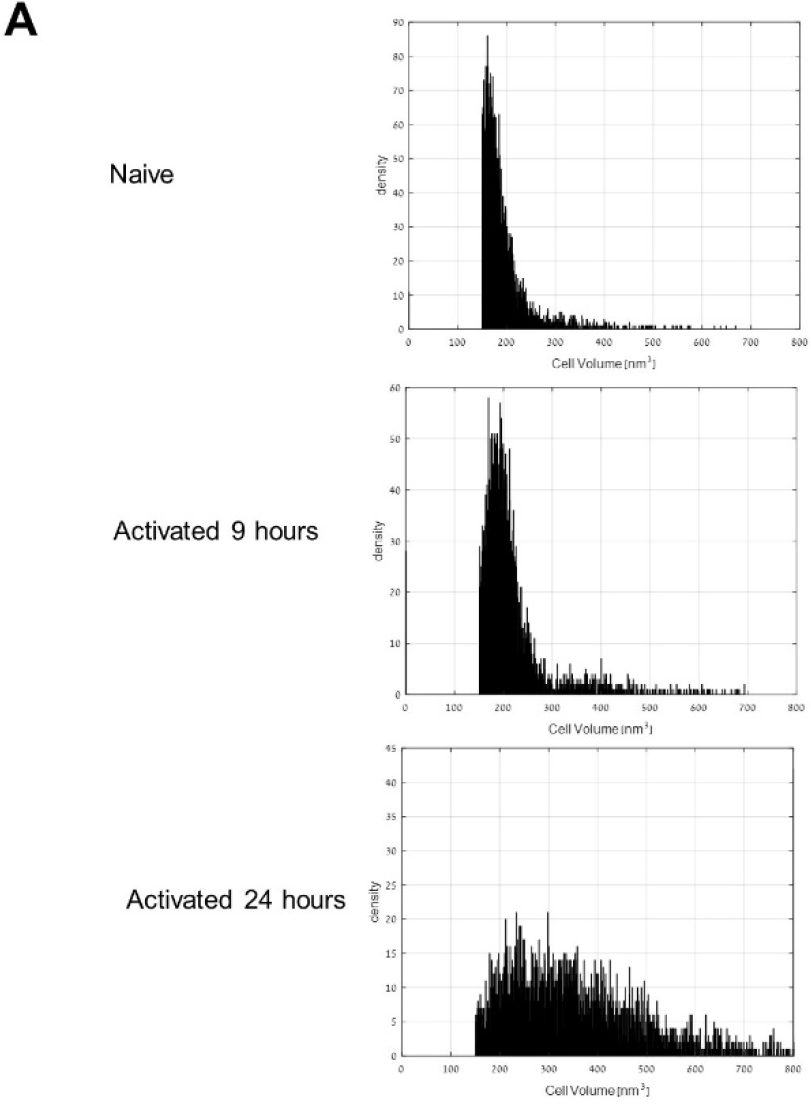
**A** Histograms showing the electronic volume of CD8+ T cells at different time points following activation.

**Figure S4:**
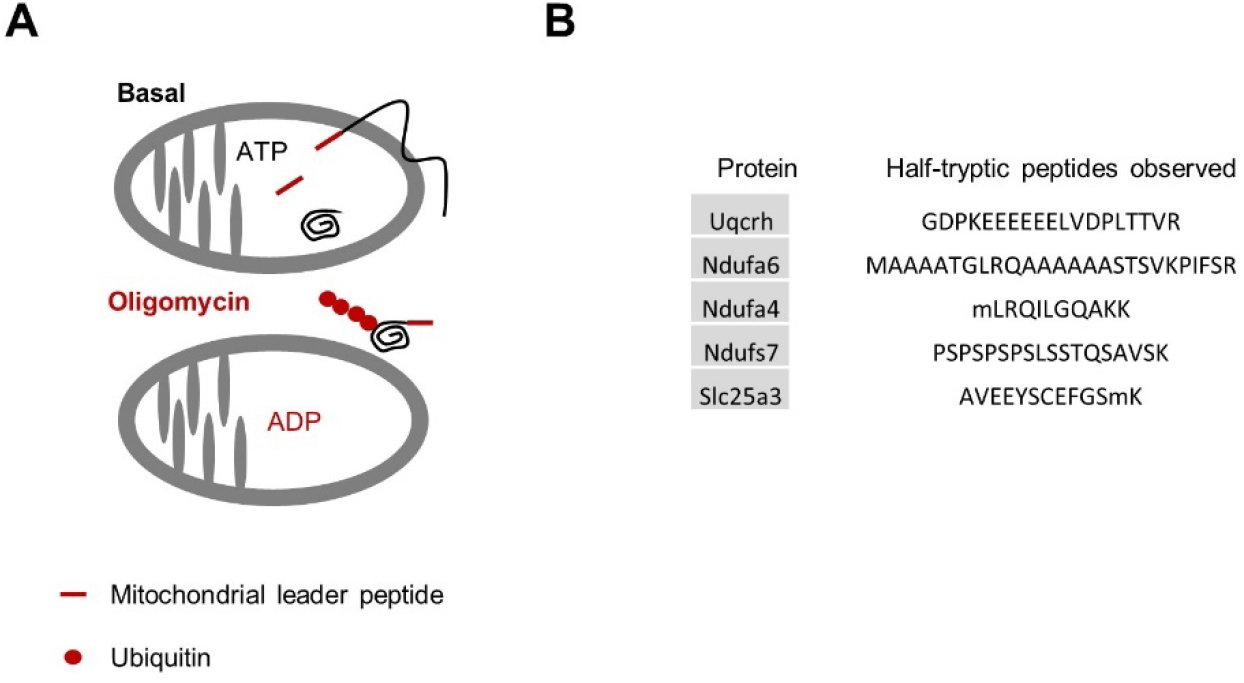
**A** Schematic of matrix ATP deficiency leading to accumulation of ubiquitinated mitochondrial matrix-localized proteins and pre-proteins. **B** Splenocytes were stimulated using anti CD3/CD28 for 9 or 12 hours. One hour prior to CD8+ T cells isolation, cells were treated with oligomycin or left untreated. Protein extract from isolated CD8+ T cells were then subjected to immunoprecipitation using anti-ubiquitin antibody following by MS analysis focusing on mitochondrial proteins and leader peptides. The table shows sequence of leader peptides and their corresponding proteins identified at 9 and 12 hours in respect to untreated samples.

**Figure S5:**
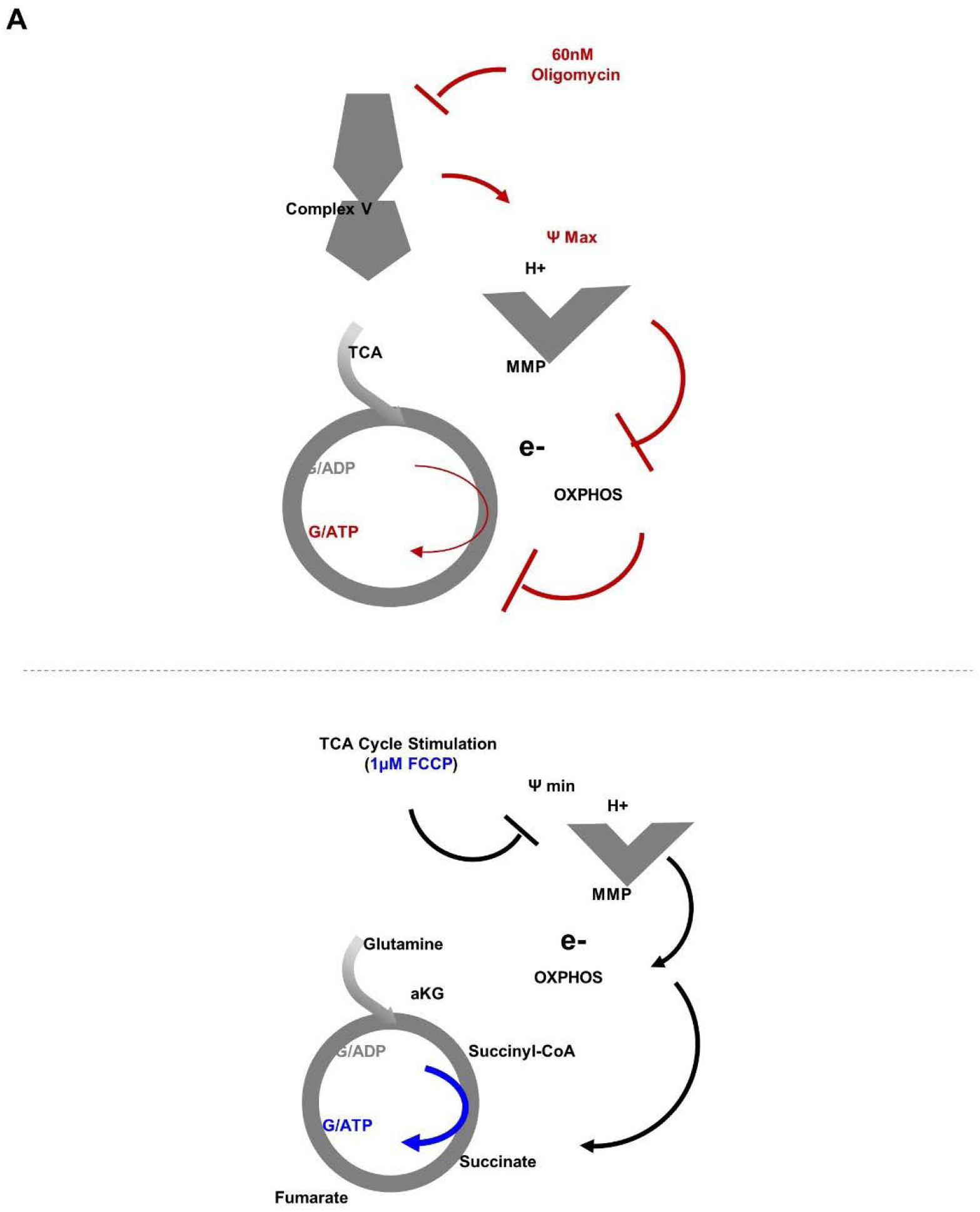
Schematic of the suggested model; respiratory restriction through inhibition of complex V by oligomycin leads to increase in mitochondrial membrane potential and decreased electron flow. Decreased electron flow leads to inhibition of TCA cycle and substrate base phosphorylation since that these processes are coupled. (TOP). Schematic of the suggested model for rescuing respiration restricted primed CD8+ T cells. FCCP treatment decreases mitochondrial membrane potential which allows both electrons and the TCA cycle to reflow, restores GTP/ATP production in the matrix by the substrate-based-phosphorylation (Bottom).

**Figure S6:**
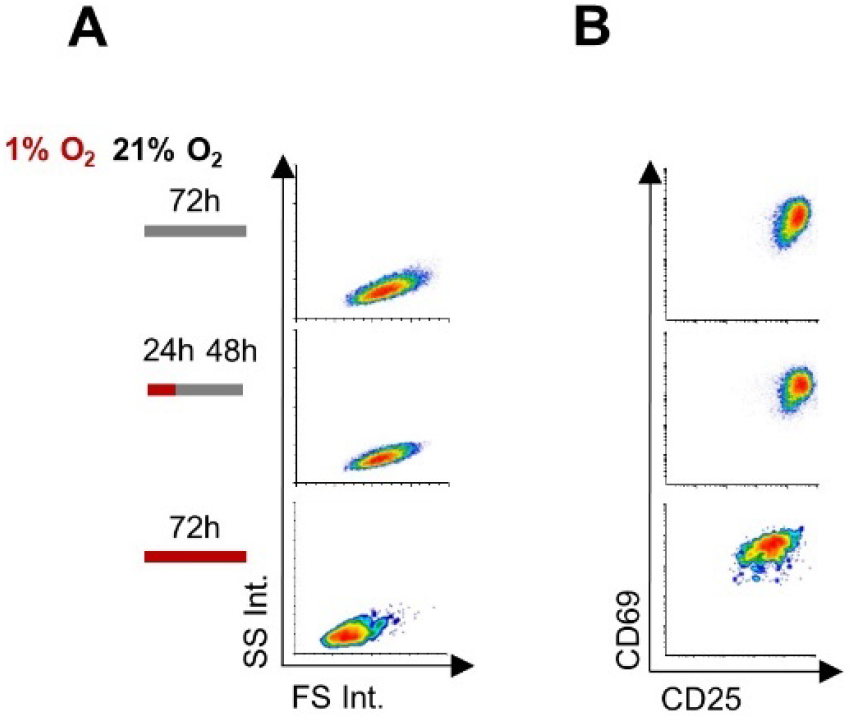
Cell Trace-labeled splenocytes were stimulated with anti-CD3/28 for 72 hours, under normal conditions (top panels), hypoxic conditions (1% O2, 5% CO2, 94% N3) for the initial 24 hours and then in normoxic condition for 48 hours (middle panels), or under ongoing hypoxic conditions (bottom panels). CD8+ T cells were then analyzed by flow cytometry for **A** size and granularity, **B** activation markers, CD25 and CD69. n= 5 biological replicates.

## SUPPLEMENTARY TABLE

**Supplementary Table 1:**
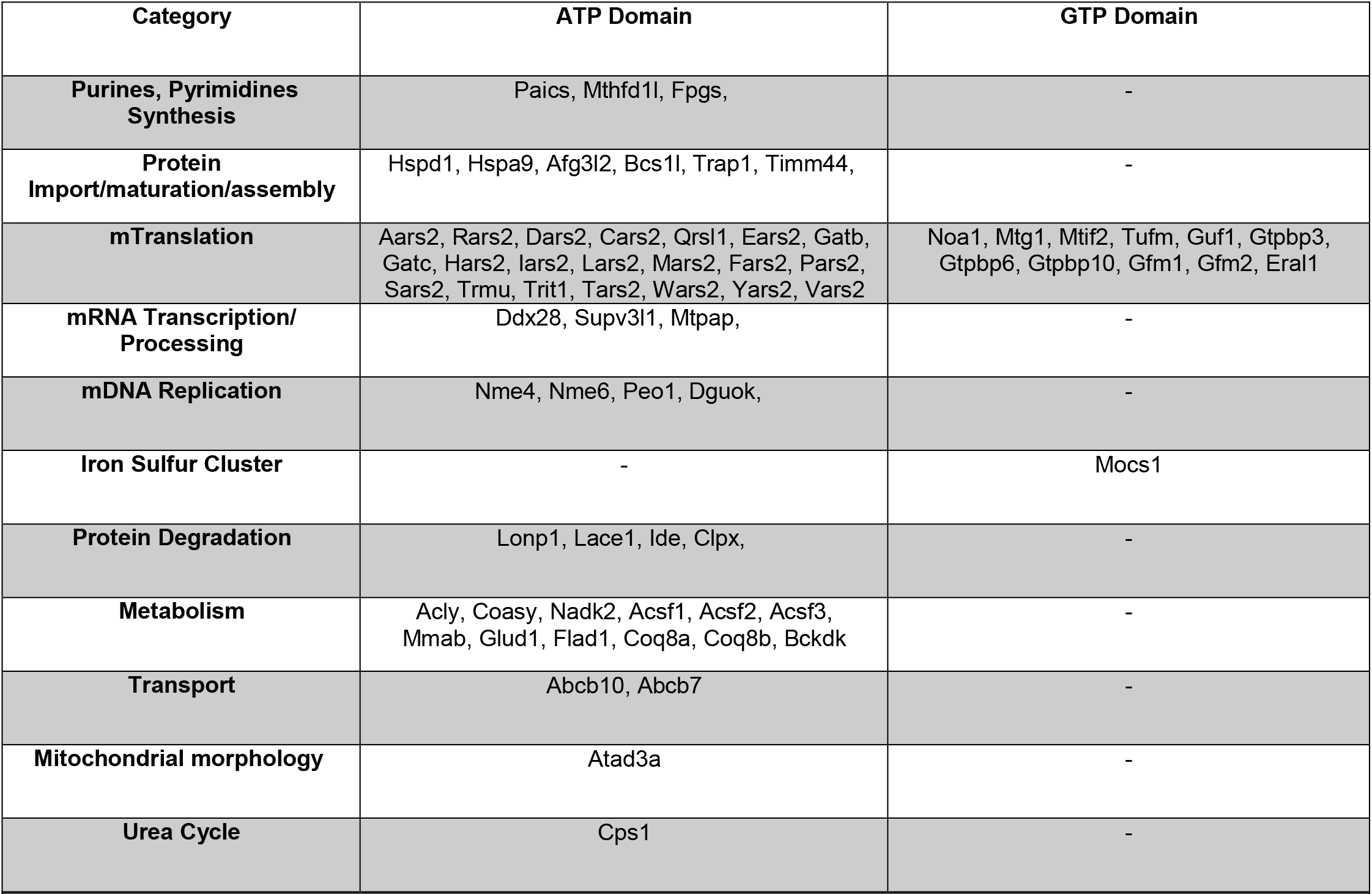
Matrix protein annotated by GO with either ATP or GTP domains

## REFERENCES

Anwar M, Kasper A, Steck AR, Schier JG. 2017. Bongkrekic Acid—a Review of a Lesser-Known Mitochondrial Toxin. Journal of Medical Toxicology. doi:10.1007/s13181-016-0577-1

Araki K, Turner AP, Shaffer VO, Gangappa S, Keller SA, Bachmann MF, Larsen CP, Ahmed R. 2009. mTOR regulates memory CD8 T-cell differentiation. Nature. doi:10.1038/nature08155

Bochud-Allemann N, Schneider A. 2002. Mitochondrial substrate level phosphorylation is essential for growth of procyclic Trypanosoma brucei. Journal of Biological Chemistry 277:32849–32854. doi:10.1074/jbc.M205776200

Buck MDD, O’Sullivan D, Klein Geltink RII, Curtis JDD, Chang CH, Sanin DEE, Qiu J, Kretz O, Braas D, van der Windt GJJW, Chen Q, Huang SCC, O’Neill CMM, Edelson BTT, Pearce EJJELL, Sesaki H, Huber TBB, Rambold ASS, Pearce EJJELL. 2016. Mitochondrial Dynamics Controls T Cell Fate through Metabolic Programming. Cell 166:63–76. doi:10.1016/j.cell.2016.05.035

Chacinska A, Koehler CM, Milenkovic D, Lithgow T, Pfanner N. 2009. Importing Mitochondrial Proteins: Machineries and Mechanisms. Cell. doi:10.1016/j.cell.2009.08.005

Chang C-H, Curtis JD, Maggi LB, Faubert B, Villarino A V., O’Sullivan D, Huang SC-C, van der Windt GJW, Blagih J, Qiu J, Weber JD, Pearce EJ, Jones RG, Pearce EL. 2013. Posttranscriptional Control of T Cell Effector Function by Aerobic Glycolysis. Cell. doi:10.1016/j.cell.2013.05.016

Chaw PS, Wen S, Wong L, Cunningham S, Campbell H, Mikolajczyk R, Nair H, Investigators R. 2019. The Journal of Infectious Diseases Acute Lower Respiratory Infections Associated With Respiratory Syncytial Virus in Children With Underlying Congenital Heart Disease: Systematic Review and Metaanalysis. The Journal of Infectious Diseases ® 1–8. doi:10.1093/infdis/jiz150

Chinopoulos C, Gerencser AA, Mandi M, Mathe K, Töröcsik B, Doczi J, Turiak L, Kiss G, Konràd C, Vajda S, Vereczki V, Oh RJ, Adam-Vizi V. 2010. Forward operation of adenine nucleotide translocase during F 0 F 1 -ATPase reversal: critical role of matrix substrate-level phosphorylation. The FASEB Journal 24:2405–2416. doi:10.1096/fj.09-149898

Cho J, Seo J, Lim CH, Yang L, Shiratsuchi T, Lee MH, Chowdhury RR, Kasahara H, Kim JS, Oh SP, Lee YJ, Terada N. 2015. Mitochondrial ATP transporter Ant2 depletion impairs erythropoiesis and B lymphopoiesis. Cell Death and Differentiation. doi:10.1038/cdd.2014.230

Cho J, Zhang Y, Park S-Y, Joseph A-M, Han C, Park H-J, Kalavalapalli S, Chun S-K, Morgan D, Kim J-S, Someya S, Mathews CE, Lee YJ, Wohlgemuth SE, Sunny NE, Lee H-Y, Choi CS, Shiratsuchi T, Oh SP, Terada N. 2017. Mitochondrial ATP transporter depletion protects mice against liver steatosis and insulin resistance. Nature Communications. doi:10.1038/ncomms14477

Doedens AL, Phan AT, Stradner MH, Fujimoto JK, Nguyen J V., Yang E, Johnson RS, Goldrath AW. 2013. Hypoxia-inducible factors enhance the effector responses of CD8 + T cells to persistent antigen. Nature Immunology. doi:10.1038/ni.2714

Gubser PM, Bantug GR, Razik L, Fischer M, Dimeloe S, Hoenger G, Durovic B, Jauch A, Hess C. 2013. Rapid effector function of memory CD8+ T cells requires an immediate-early glycolytic switch. Nature Immunology 14:1064–1072. doi:10.1038/ni.2687

Kaskinen AK, Helve O, Andersson S, Kirjavainen T, Martelius L, Mattila IP, Rautiainen P, Pitkänen OM. 2016. Chronic Hypoxemia in Children With Congenital Heart Defect Impairs Airway Epithelial Sodium Transport. Pediatric Critical Care Medicine 17:45–52. doi:10.1097/PCC.0000000000000568

Kent BD, Mitchell PD, Mcnicholas WT. 2011. Hypoxemia in patients with COPD: Cause, effects, and disease progression. International Journal of COPD. doi:10.2147/COPD.S10611

Kherad O, Kaiser L, Bridevaux PO, Sarasin F, Thomas Y, Janssens JP, Rutschmann OT. 2010. Upper-respiratory viral infection, biomarkers, and COPD exacerbations. Chest 138:896–904. doi:10.1378/chest.09-2225

Lee O, O’Brien PJ. 2010. Modifications of Mitochondrial Function by ToxicantsComprehensive Toxicology: Second Edition. Elsevier Inc. pp. 411–445. doi:10.1016/B978-0-08-046884-6.00119-6

Lunt SY, Vander Heiden MG. 2011. Aerobic Glycolysis: Meeting the Metabolic Requirements of Cell Proliferation. Annual Review of Cell and Developmental Biology 27:441–464. doi:10.1146/annurev-cellbio-092910-154237

MacIver NJ, Michalek RD, Rathmell JC. 2013. Metabolic regulation of T lymphocytes. Annu Rev Immunol. doi:10.1146/annurev-immunol-032712-095956

MacKay GM, Zheng L, Van Den Broek NJF, Gottlieb E. 2015. Analysis of Cell Metabolism Using LC-MS and Isotope TracersMethods in Enzymology. doi:10.1016/bs.mie.2015.05.016

Martínez-Reyes I, Diebold LP, Kong H, Zhao Y, Deberardinis RJ, Chandel Correspondence NS. 2016. TCA Cycle and Mitochondrial Membrane Potential Are Necessary for Diverse Biological Functions. Molecular Cell 61. doi:10.1016/j.molcel.2015.12.002

O’Brien P, Smith PA. 1994. Chronic hypoxemia in children with cyanotic heart disease. Critical care nursing clinics of North America 6:215–26.

Pearce EL, Walsh MC, Cejas PJ, Harms GM, Shen H, Wang LS, Jones RG, Choi Y. 2009. Enhancing CD8 T-cell memory by modulating fatty acid metabolism. Nature 460:103–107. doi:10.1038/nature08097

Pfanner N, Warscheid B, Wiedemann N. 2019. Mitochondrial proteins: from biogenesis to functional networks. Nature Reviews Molecular Cell Biology. doi:10.1038/s41580-018-0092-0

Pham AH, Mccaffery JM, Chan DC. 2012. Mouse lines with photo-activatable mitochondria to study mitochondrial dynamics. Genesis. doi:10.1002/dvg.22050

Phan AT, Doedens AL, Palazon A, Tyrakis PA, Cheung KP, Johnson RS, Goldrath AW. 2016. Constitutive Glycolytic Metabolism Supports CD8+T Cell Effector Memory Differentiation during Viral Infection. Immunity 45:1024–1037. doi:10.1016/j.immuni.2016.10.017

Phan AT, Goldrath AW. 2015. Hypoxia-inducible factors regulate T cell metabolism and function. Molecular Immunology. doi:10.1016/j.molimm.2015.08.004

Rackham O, Busch JD, Matic S, Siira SJ, Kuznetsova I, Atanassov I, Ermer JA, Shearwood AMJ, Richman TR, Stewart JB, Mourier A, Milenkovic D, Larsson NG, Filipovska A. 2016. Hierarchical RNA Processing Is Required for Mitochondrial Ribosome Assembly. Cell Reports 16:1874–1890. doi:10.1016/j.celrep.2016.07.031

Rambold AS, Pearce EL. 2017. Mitochondrial Dynamics at the Interface of Immune Cell Metabolism and Function. Trends in Immunology. doi:10.1016/j.it.2017.08.006

Rappsilber J, Mann M, Ishihama Y. 2007. Protocol for micro-purification, enrichment, pre-fractionation and storage of peptides for proteomics using StageTips. Nature Protocols. doi:10.1038/nprot.2007.261

Ron-Harel N, Santos D, Ghergurovich JM, Sage PT, Reddy A, Lovitch SB, Dephoure N, Satterstrom FK, Sheffer M, Spinelli JB, Gygi S, Rabinowitz JD, Sharpe AH, Haigis MC. 2016. Mitochondrial Biogenesis and Proteome Remodeling Promote One-Carbon Metabolism for T Cell Activation. Cell Metabolism 24:104–117. doi:10.1016/j.cmet.2016.06.007

Schwimmer C, Lefebvre-Legendre L, Rak M, Devin A, Slonimski PP, Di Rago JP, Rigoulet M. 2005. Increasing mitochondrial substrate-level phosphorylation can rescue respiratory growth of an ATP synthase-deficient yeast. Journal of Biological Chemistry 280:30751–30759. doi:10.1074/jbc.M501831200

Sgarbi G, Barbato S, Costanzini A, Solaini G, Baracca A. 2018. The role of the ATPase inhibitor factor 1 (IF1) in cancer cells adaptation to hypoxia and anoxia. Biochimica et Biophysica Acta - Bioenergetics 1859:99–109. doi:10.1016/j.bbabio.2017.10.007

Solaini G, Baracca A, Lenaz G, Sgarbi G. 2010. Hypoxia and mitochondrial oxidative metabolism. Biochimica et Biophysica Acta - Bioenergetics. doi:10.1016/j.bbabio.2010.02.011

Tu YT, Barrientos A. 2015. The Human Mitochondrial DEAD-Box Protein DDX28 Resides in RNA Granules and Functions in Mitoribosome Assembly. Cell Reports 10:854–864. doi:10.1016/j.celrep.2015.01.033

Van Der Windt GJW, O’Sullivan D, Everts B, Huang SCC, Buck MD, Curtis JD, Chang CH, Smith AM, Ai T, Faubert B, Jones RG, Pearce EJ, Pearce EL. 2013. CD8 memory T cells have a bioenergetic advantage that underlies their rapid recall ability. Proceedings of the National Academy of Sciences of the United States of America 110:14336–14341. doi:10.1073/pnas.1221740110

Wang G, Chen HW, Oktay Y, Zhang J, Allen EL, Smith GM, Fan KC, Hong JS, French SW, McCaffery JM, Lightowlers RN, Morse HC, Koehler CM, Teitell MA. 2010. PNPASE regulates RNA import into mitochondria. Cell. doi:10.1016/j.cell.2010.06.035

Wiedemann N, Pfanner N. 2017. Mitochondrial Machineries for Protein Import and Assembly. Annual Review of Biochemistry 86:685–714. doi:10.1146/annurev-biochem-060815-014352

Yu AY, Shimoda LA, Iyer N V., Huso DL, Sun X, McWilliams R, Beaty T, Sham JSK, Wiener CM, Sylvester JT, Semenza GL. 1999. Impaired physiological responses to chronic hypoxia in mice partially deficient for hypoxia-inducible factor 1α. Journal of Clinical Investigation 103:691–696. doi: 10.1172/JCI5912

